# An adhesion GPCR regulates cell adhesion and mating in the closest living relatives of metazoans

**DOI:** 10.64898/2026.06.27.734982

**Authors:** Alain Garcia De Las Bayonas, Stefany Gonzalez, Nicole King

## Abstract

The transition to metazoan multicellularity required the evolution of cell-surface receptors that coordinate adhesion and signaling under changing environmental conditions. We investigated potential regulators of cell interactions in the choanoflagellate *Salpingoeca rosetta*, one of the closest living relatives of metazoans. Here, we identify Cupidon, an adhesion G protein-coupled receptor that acts as a dual-function regulator of cell adhesion and mating. Under well-fed (i.e., nutrient-replete) conditions, Cupidon suppresses cell aggregation by inhibiting N-acetylglucosamine-dependent collar-mediated adhesion. Starvation of *S. rosetta* triggers gametogenesis, resulting in anisogametes: female gametes with an elongated collar and male gametes that form a basal protrusion, the “fertilopod.” Cupidon undergoes concurrent changes in proteolytic processing and localization, ultimately concentrating at the gamete contact interface, where it promotes gamete fusion. Together, our findings reveal that aGPCR-mediated regulation of cell adhesion predates the origin of metazoans.

## INTRODUCTION

Choanoflagellates, the closest living relatives of metazoans^1–3^, offer a powerful system for investigating the evolution and regulation of cell-state transitions. The evolutionary relationship between choanoflagellates and metazoans is reflected in the conservation of gene families that underlie metazoan cell biology, including those encoding signaling receptors, adhesion proteins, and cytoskeletal regulators^3–6^. In addition, like sponge choanocytes and certain cells in most metazoans, choanoflagellates possess a collar complex, a distinct cellular module of actin-filled microvilli surrounding an apical flagellum^7–9^. In choanoflagellates and sponge choanocytes, the undulating flagellum generates water currents that concentrate bacterial prey on the outer surface of the collar. Finally, choanoflagellates can form multicellular structures, some of which resemble the organization of metazoan tissues^10–13^.

The choanoflagellate *Salpingoeca rosetta* is experimentally tractable^14–16^ and exhibits a flexible life history: depending on environmental conditions, it can exist as solitary flagellated cells, amoeboid cells, thecate cells, or can transition into multicellular forms (chains or rosette colonies)^17^. Under nutrient-replete conditions, chains form by incomplete cytokinesis, while rosette colonies develop clonally from a single cell in response to bacterial lipid cues ^18,19^. *S. rosetta* also undergoes sexual reproduction, which can be induced either by starvation or treatment with EroS, a chondroitinase secreted by *Vibrio fischeri* and related bacteria^20^. Mating involves sequential steps — cell aggregation, gamete differentiation, pairing, and fusion — each of requiring coordinated changes in cell adhesion and signaling, the molecular underpinnings of which remain poorly understood.

G protein-coupled receptors (GPCRs) are prime candidates for regulating *S. rosetta* cell differentiation and behavior. Conserved across eukaryotes^21,22^, GPCRs respond to diverse extracellular cues, including lipids, amino acids, and microbial metabolites^23–25^. GPCRs regulate cell-state transitions ranging from fungal mating to vertebrate gametogenesis^26–28^. For example, nutrient availability can be sensed through GPCRs: in fungi, glucose-and nitrogen-sensing GPCRs directly couple nutritional status to cell-state transitions, including the decision to enter meiosis and mate^29,30^.

A specialized subfamily — the adhesion GPCRs (aGPCRs) — is defined by an extracellular region linked to a seven-transmembrane-helix (7TM) signaling module by a GPCR-autoproteolysis-inducing (GAIN) domain ^31,32^. A subset of aGPCRs that harbor large extracellular regions containing adhesion-related protein domains (hereafter, multidomain aGPCRs) is restricted to filastereans, choanoflagellates, and metazoans ^22^. In metazoans, multidomain aGPCRs are central regulators of cell-cell and cell-matrix adhesion^32–34^. In *S. rosetta*, multidomain aGPCRs constitute the largest GPCR subfamily, representing 12 of the 23 encoded GPCRs^22^. The combination of adhesion-competent extracellular domains with the 7TM signaling module allows aGPCRs to couple cell adhesion to intracellular signal transduction^34^. Yet, to date, no choanoflagellate GPCR has been functionally characterized.

Here, we identify Cupidon, a choanoflagellate-specific aGPCR, as a dual-function regulator of cell adhesion and mating in *S. rosetta*. Our findings demonstrate that aGPCR-mediated control of cell–cell adhesion predates the origin of metazoans.

## RESULTS

### The adhesion GPCR Cupidon suppresses cell aggregation under nutrient-replete culture conditions

*S. rosetta* expresses 12 aGPCRs, including one, hereafter referred to as *cupidon* (French for Cupid, the mischievous god of love), that was intriguing because of its conservation in a clade of nine out of the 23 choanoflagellates analyzed (Figure 1A). Cupidon likely evolved in choanoflagellates, as no orthologs were detected in filastereans or metazoans (Figure 1A). Multiple sequence alignment of Cupidon from all 9 choanoflagellates revealed the conservation of 13 protein motifs and domains (Figure 1B and Figure S1A and B, File S2), including a Hormone Receptor Motif (HRM), a GPCR autoproteolysis-inducing (GAIN) domain characteristic of aGPCRs^31,35^, and the diagnostic seven transmembrane helices (7TMs) of GPCRs. The extracellular portion (or N-terminal fragment) also contained three domains that are commonly involved in cell-cell and cell-matrix interactions: Epidermal Growth Factor (EGF) domains, a Concanavalin A-like lectin domain (ConA), and a Thrombospondin type-1 (TSP1) domain^36–41^. We also found a previously uncharacterized Mystery Domain (MD) that is predicted to adopt a beta-strand-rich fold (Figure S1D) that is conserved in diverse non-GPCR proteins from bacteria, viruses, and eukaryotes (Alveolates, Haptista, and Parmales^42^; Figure S1C, E, and Video S1). The C-terminal fragment of Cupidon contained no readily-identifiable domains, although the SYF motif at the C-terminal end of Cupidon from *S. rosetta* and *Microstomoeca roanoka* (Figure S1F) may interact with Class I PDZ domains^43–45^ as it does in some metazoan aGPCRs^32^.

**Figure 1.**
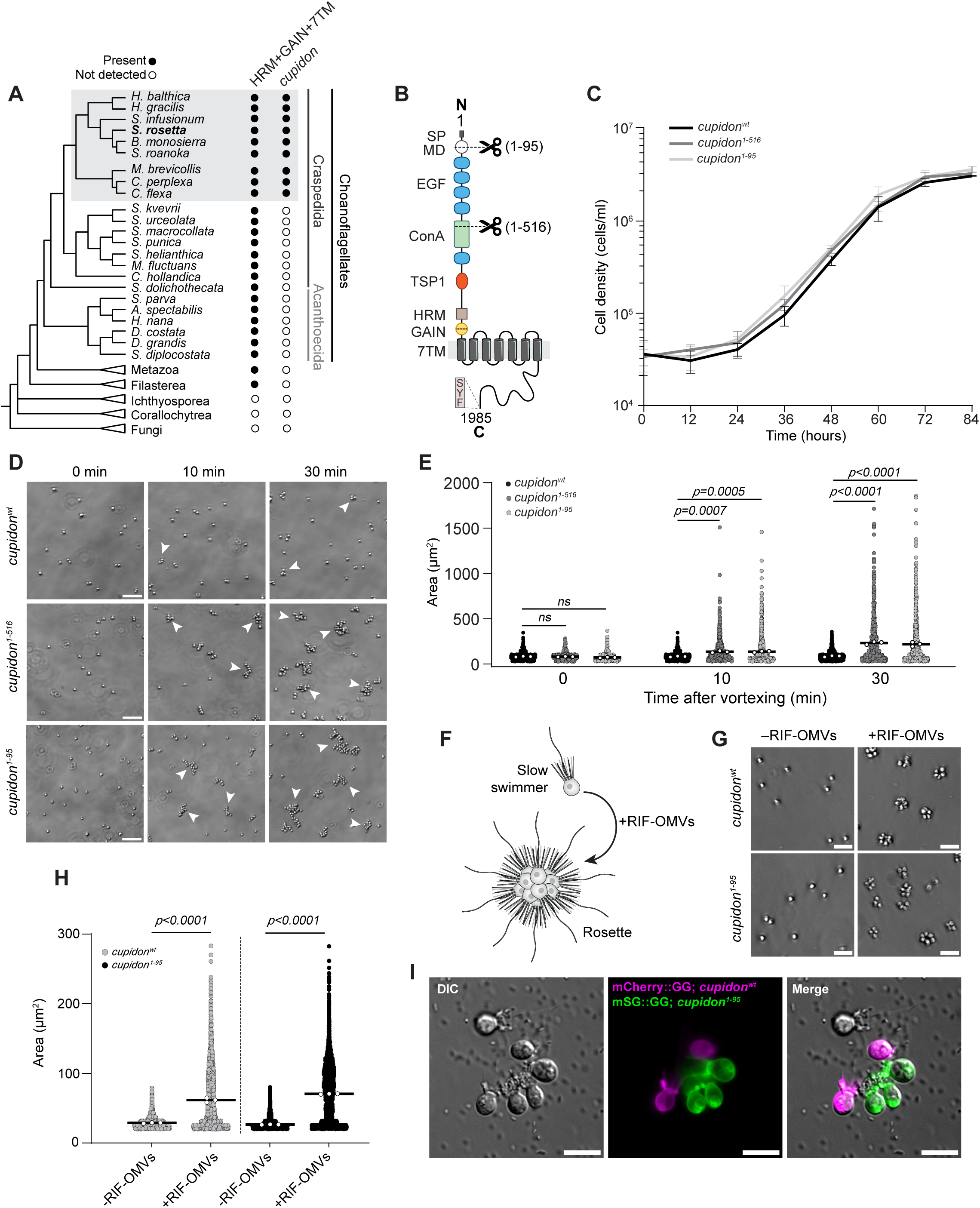
Cupidon inhibits cell aggregation in cultures grown under replete conditions. (A) Multidomain aGPCRs originated in the last common ancestor of filastereans, choanoflagellates, and metazoans^22^. The *cupidon* gene is restricted to a subset of nine choanoflagellate species within craspedids (grey box). (B) Schematic of the protein domain architecture of Cupidon and edits that generated the truncated protein variants (Cupidon^1–95^ and Cupidon^1–516^) examined in this study. Cupidon contains a signal peptide (SP, grey rectangle), a Mystery domain (MD, white circle), five EGF domains (blue ovals), a ConA domain (light green rectangle), a TSP1 domain (orange circle), an HRM domain (light brown square), a GAIN domain (yellow circle), a 7TM domain (grey cylinders), and a putative Class I PDZ binding motif (SYF) at the carboxyl end of the receptor. (C) Disruption of *cupidon* in the *cupidon^1–516^* (dark grey line) and *cupidon*^1–95^ (light grey line) strains did not affect cell proliferation compared with wild-type cells (black line). Three biological replicates were analyzed per experimental condition; the mean values were plotted, with standard deviations shown as error bars. (D) Representative time series showing ectopic cell aggregation in replete-grown *cupidon^1–516^* and *cupidon*^1–95^ mutant cultures compared with *cupidon*^wt^ cells, 0, 10, and 30 min. after vortexing. Examples of cell clumps are depicted with white arrows; while a few small clumps can be found in *cupidon*^wt^ cultures, their size and number are greatly increased in *cupidon* mutants. Scale bars = 50 μm. (E) Quantification of cell clump area from experiments performed as in (D). Data are presented as scatter plots, each dot corresponding to the projected area of an object (single cells or clumps). Three biological replicates were analyzed for each experimental condition; white dots represent means per replicate; bars denote means across replicates. A minimum of 923 clump areas were plotted for each condition. P-values were calculated from a one-way ANOVA performed independently at each time point. (F) Schematic of RIF-OMV-induced transition to rosette development. (G) Cupidon^1–95^ mutant cells retained their ability to form rosettes upon induction with outer membrane vesicles from *Algoriphagus machipongonensis* bacteria (RIF-OMVs). Representative DIC images of a 48-hour incubation with or without RIF-OMVs followed by vortexing. While vortexed uninduced cultures (-RIF-OMVs) from both genotypes contained only single cells, shear-resistant rosettes formed and persisted in both genotypes upon RIF-OMV treatment (+RIF-OMVs), followed by vortexing. Scale bars = 20 μm. See Figure S4D. (H) Quantification of rosette induction in wild-type and *cupidon*^1–95^ mutant cultures, as shown in (G). Scatter plots show the projected area of each object (single cell or rosette); three biological replicates were analyzed per experimental condition; white dots represent means per replicate; bars denote means across replicates. A minimum of 4736 single cell or rosette areas were plotted per condition. P-values were calculated from a two-way ANOVA with Tukey’s multiple comparisons test. (I) Cupidon^1–95^ cells adhere to wild-type cells. Wild-type cells expressing the fluorescent membrane marker mCherry::GG were mixed with *cupidon*^1–95^ mutant cells expressing the fluorescent membrane marker mStayGold::GG. The resulting culture was vortexed and imaged after 30 min. Mutant cells adhered to each other and to wild-type cells. See also Video S5. Scale bar = 10 μm.

To investigate the function of Cupidon, we used CRISPR-mediated genome editing to introduce early stops at codons 95 (*cupidon*^1–95^) and 516 (*cupidon^1–516^*). The edited alleles of *cupidon* encode truncated proteins that lack the C-terminal cytoplasmic tail, the transmembrane domains, and most of the N-terminal fragment (Figure 1B and S2A). While the *cupidon^1–516^* allele retained the uncharacterized MD, four EGF domains, and part of the ConA domain, the truncated *cupidon*^1–95^ protein preserved only a portion of the MD domain.

The resulting *S. rosetta* mutant lines were grown and analyzed under standard culture conditions (hereafter referred to as “nutrient replete”) that support the proliferation of prey bacteria and *S. rosetta*. Neither mutant culture exhibited a growth defect relative to the wild type (Figure 1C), suggesting that *cupidon* is not essential for growth, cell division, or prey capture.

However, both mutant lines formed large clumps of cells in culture (Figure 1D-E, S2B-C; Video S2). Unlike rosettes, in which all cells share the same polarity and the rosette structure is resistant to shear stress^9,11,17,46,47^, clumps of *cupidon*^1–95^ and *cupidon^1–516^* cells appeared disorganized and readily separated into solitary cells upon exposure to shear. Moreover, *cupidon* mutant cells reaggregated into new clumps within 10 min after vortexing (Figures 1D-1E, S2C; Video S3), in contrast to rosette development, which requires cell division and takes at least 16 hours^11,47^. Because both *cupidon^1–516^* and *cupidon*^1–95^ lines displayed the same phenotype, we focused the remainder of our analyses on *cupidon*^1–95^, as it represents a more complete truncation.

A previous genetic screen identified *S. rosetta* mutants that formed clumps and failed to develop into rosettes^46,48^. We therefore asked whether *cupidon* mutants likewise lost the ability to produce rosettes. Treatment of wild-type and *cupidon*^1–95^ cells with rosette-inducing factors from *Algoriphagus machipongonensis* bacteria^18,19,49^ (RIF-OMVs) caused both genotypes to develop into morphologically indistinguishable rosettes (Figure 1F-H). Nonetheless, undisturbed *cupidon*^1–95^ rosettes adhered to each other, akin to the clumping behavior of *cupidon*^1–95^ solitary swimmers, while wild-type rosettes remained separated (Figure S2D, Video S4). Rosettes of *cupidon*^1–95^ cells maintained their structural integrity under shear stress (Figure 1G, S2D), demonstrating that *cupidon*^1–95^ mutant cells develop into normal rosettes. Thus, in contrast to inferences made from analyses of the clumping mutants Jumble and Couscous^46^, cell aggregation and clonal multicellularity are controlled by distinct pathways.

Importantly, we found that *cupidon*^1–95^ cells aggregated with wild-type cells (Figure 1I and Video S5), demonstrating that the adhesive interface does not require homophilic interactions between mutant cells. In addition, wild-type cells incubated with conditioned medium from *cupidon*^1–95^ cultures did not form clumps, arguing against the secretion of a pro-adhesive factor by the mutant (Figure S2E-F). Together, these data suggest that cells are naturally adherent under replete conditions and that Cupidon acts cell-autonomously to suppress cell-cell adhesion in this context, as revealed by the ectopic aggregation observed in *cupidon* mutants.

### Aggregation by *cupidon* mutants resembles starvation-induced aggregation

While *S. rosetta* reproduces mitotically when grown under replete culture conditions^50^ (Figure 2A, B), cultures induced to mate form transient clumps of haploid cells before pairing and fusion (Figure 2A, B; Video S6). Two mating triggers have been identified: (1) starvation ^50^ (Figure 2A), and (2) treatment with a secreted bacterial chondroitinase, EroS, produced by *Vibrio fischeri* and related bacteria^20^ (Figure 2A). After prolonged culturing in spent medium, asexual swimmers transition into a mixed population of morphologically distinct male and female gametes^50^ (Figure 2A). In contrast, mating gametes show no signs of differentiation in *EroS*-treated cultures compared with asexual cells^20^ (Figure 2A and 2B). We investigated whether cell aggregation in Cupidon mutants is associated with either mating pathway.

**Figure 2.**
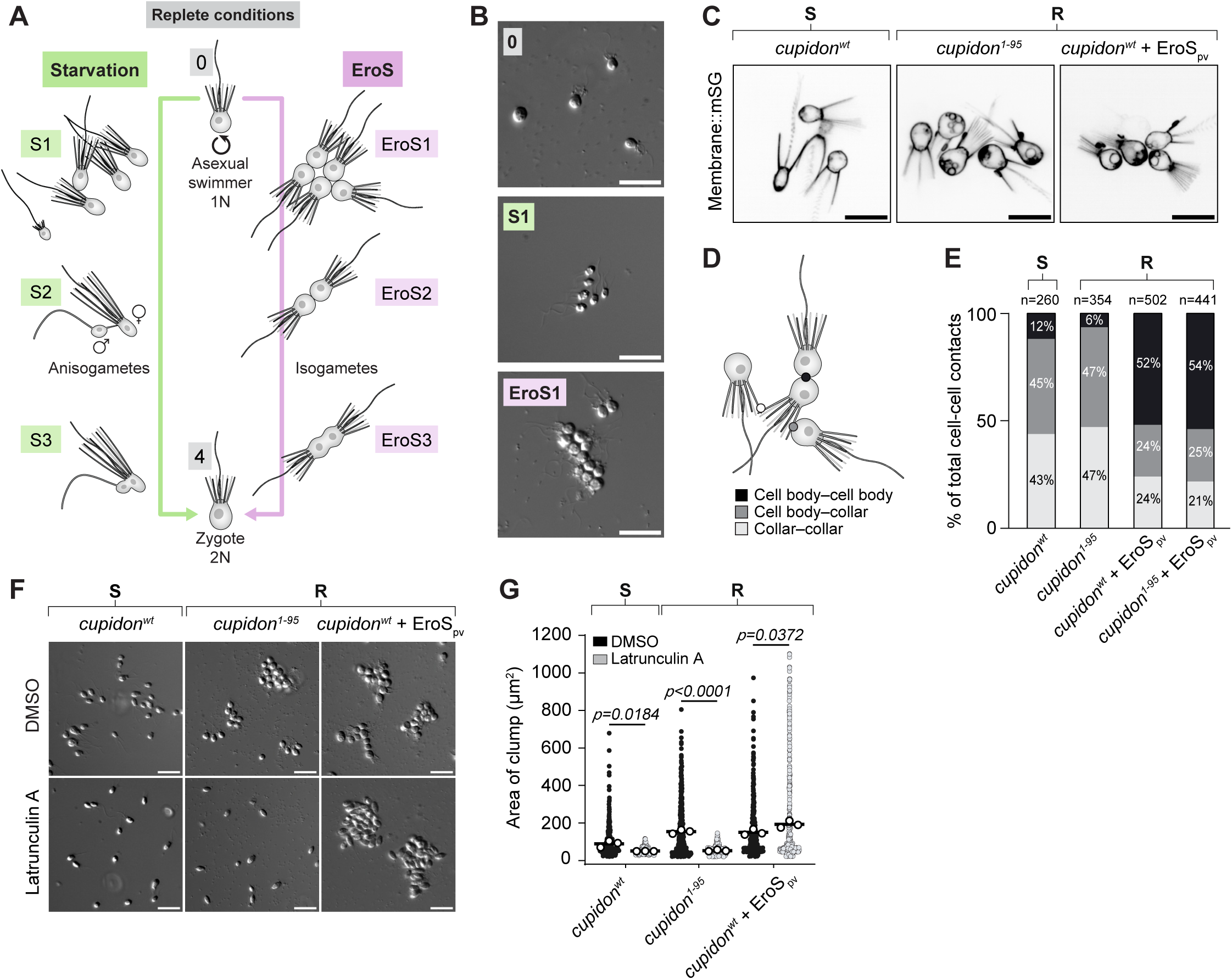
**Cupidon**^1–95^ **phenocopies starvation-induced cell aggregation.** (A) Mating can be induced by starvation^50^ (left pathway) or by treatment with the EroS protein^20^ (right pathway). Both involve clumping of haploid cells (S1 and EroS1, respectively) prior to pairing (S2, EroS2), gamete fusion (S3, EroS3), and nuclear fusion leading to the 2N zygote (4). The starvation pathway leads to the production of two distinct gametes, which differ in size and form (anisogametes) while EroS induction results in pairing of isogametes (EroS2). (B) Representative images of asexual cells grown in replete conditions (0), a clump of cells formed after maintenance in spent medium (S1), and a clump of cells in replete medium after treatment with EroS protein (EroS1). Scale bars = 20 μm. Also see Video S6. (C) Imaging of cells expressing the membrane marker mStayGold::GG (mStayGold fused to a *S. rosetta* geranylgeranylation sequence) reveals the spatial organization of cells and the geometry of cell-cell contacts within clumps. Scale bars = 10 μm. (D) Cell-cell interactions fall into three categories: cell body-cell body interactions (black), cell body-collar interactions (dark grey), and collar-collar interactions (light grey). (E) Cell-cell contacts in *cupidon*^1–95^ cells predominantly involve the collar and most closely match those of *cupidon^wt^* cells maintained in spent medium. EroS_pv_-treated *cupidon^wt^* cells, in contrast, interact primarily through cell body–cell body interactions. Treatment of *cupidon*^1–95^ cells with EroS_pv_ resulted in a similar abundance of cell body–cell body interactions. The numbers above the bars indicate the number of clumps analyzed per experimental condition. Two biological replicates were analyzed per condition. (F and G) Actin depolymerization with Latrunculin A disrupted the actin-based collar (also see Video S7), thereby preventing clumping in starved *cupidon^wt^* cells and replete *cupidon*^1–95^ cells, but did not affect clumping in EroS_pv_-treated cells. Scale bars = 20 μm. For Figure 2G, three biological replicates were analyzed per condition. A minimum of 498 single-cell or clump areas were plotted per condition; white dots represent means per replicate; bars denote means across replicates. P-values were calculated from a two-tailed t-test performed independently for each experimental condition.

We compared the mode of cell-cell binding within cell aggregates formed by *cupidon*^1–95^ mutant cells grown in replete conditions with that of aggregates formed by either starved *cupidon*^wt^ cells or *cupidon*^wt^ cultures treated with EroS commercially prepared from *Proteus vulgaris* bacteria (EroS_pv_)^20,49^. Starved wild-type cells and *cupidon*^1–95^ cells grown in replete conditions predominantly aggregated through collar-collar (∼45% and ∼47%, respectively) and cell body-collar interactions (∼43% and ∼47%, respectively; Figure 2C-E), while cells in EroS_pv_-induced mating clumps interacted primarily through cell body-cell body contacts (∼52%; Figures 2C-E; Video S7). Treating *cupidon*^1–95^ cells with EroS_pv_ shifted the mode of interaction to favor cell body-cell body contacts (∼54%), thus more closely resembling clumps from EroS_pv_-induced mating cells (Figure 2E, S3D). These results revealed two modes of cell–cell adhesion in response to different mating cues in *S. rosetta*. They also suggested that *cupidon*^1–95^ cells and starved *cupidon*^wt^ cells aggregate through a common mechanism.

Because the collar appeared to be the preferred adhesive interface in both *cupidon*^1–95^ and wild-type cells grown in spent media, we tested whether the cell collar was necessary to sustain clumping in both conditions. We treated *cupidon*^1–95^ cells grown in replete media and starved *cupidon*^wt^ cells with Latrunculin A, a potent inhibitor of F-actin assembly that results in the loss of the actin-rich collar without affecting cell motility (Video S7). Latrunculin A treatment completely abolished cell aggregation in both *cupidon*^1–95^ and starved *cupidon*^wt^ cells (Figures 2F and 2G), while showing no effect on clumping in EroS_pv_-treated cells. These data confirm the necessity of the collar for clump formation in both starved *cupidon*^wt^ cells and replete-grown *cupidon*^1–95^ cells, suggesting that Cupidon is required to repress collar adhesion in *S. rosetta* cells grown under replete conditions.

### Starvation alters Cupidon processing and localization in gametes

We hypothesized that changes in transcription, translation, or the post-translational processing of *cupidon* could contribute to aggregation and mating during starvation. The transcription of *cupidon* increases over the course of starvation, reaching a ∼2.3-3X upregulation in mating cultures relative to asexual cells (Figure S4A and Table S1), revealing that any reduction in the anti-adhesive function of Cupidon cannot be explained by transcriptional repression. Thus, we inferred that any change in Cupidon activity between asexual and sexual cells might be regulated post-transcriptionally. To assess the biochemical regulation and localization of Cupidon, we engineered two endogenously tagged knock-in lines of Cupidon to introduce either an ALFA epitope^51^ or a monomeric StayGold fluorescent reporter^52^ in the carboxy-terminal region (Figure 3A, S4B-D). Importantly, neither of the tagged lines formed clumps under replete conditions, suggesting that the two in-frame tag insertions preserved Cupidon function (Figure S4E-F; Video S8).

**Figure 3.**
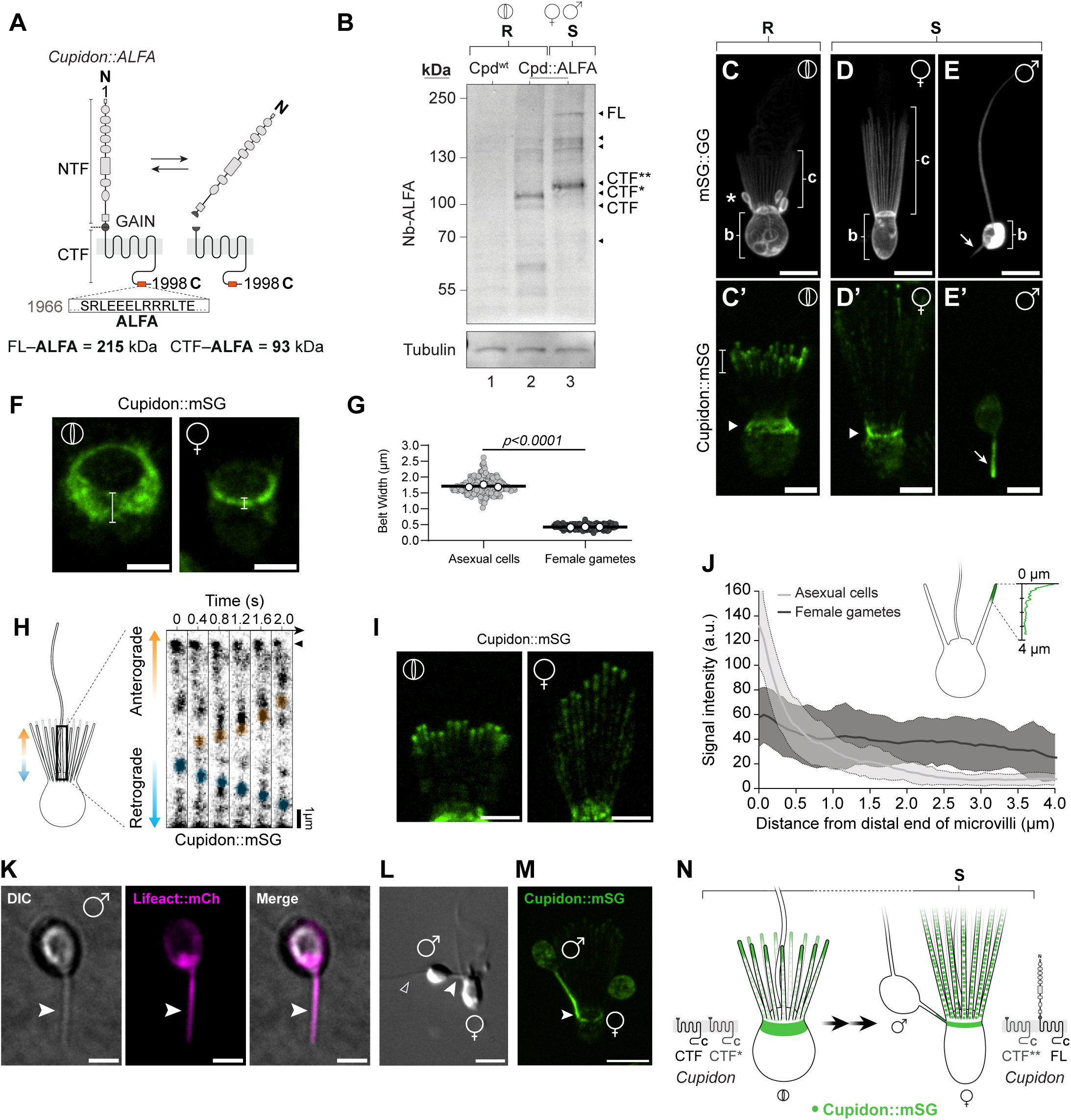
Alternative processing and relocalization of Cupidon during gametogenesis. (A) Endogenous tagging of Cupidon through the insertion of an ALFA tag sequence (shown) at position 1966. The ALFA tag allows detection of full-length Cupidon (FL-ALFA; 215 kDa) prior to cleavage and the C-terminal fragment (CTF-ALFA; 93 kDa) by using an ALFA-specific nanobody. (B) Western blot (Nb-ALFA) reveals that Cupidon undergoes differential post-translational processing in cells grown in replete (R) or spent (S) medium. The ALFA-specific nanobody detected no background signal in wild-type cells (Cpd^wt^) grown in replete conditions (lane 1). In Cpd::ALFA cells grown in replete conditions, the ALFA-specific nanobody detected a band at the predicted size of the C-terminal fragment after cleavage at the GAIN domain (CTF), a more prominently band of slightly larger apparent molecular weight (∼110 kDa; CTF*), and additional faint bands at ∼130–150 kDa and ∼60 kDa (lane 2). In Cpd::ALFA cells maintained in spent medium, the nanobody detected bands at ∼ 220 kDa, corresponding to full-length Cupidon (FL) and a dark band at ∼115 kDa (CTF**), along with additional faint bands between ∼130–150 kDa (lane 3). Tubulin is shown as a loading control. (C-E’) Differences in cell architecture and Cupidon localization in asexual cells ⦶ (C, C’), female gametes **♀** (D, D’), and male gametes ♂ (E, E’). *S. rosetta* cells were edited to express either mStayGold::GG, which allowed detection of the cell membrane (C, D, E), or Cupidon::mStayGold, which allowed monitoring of Cupidon localization in live cells (C’, D’, E’). (C, D, E) mStayGold::GG revealed that gametogenesis in spent medium (S) results in dramatic reorganization of cell architecture. (C) Asexual cells have a rounded cell body (b), an apical collar of microvilli (c), and an apical flagellum (not visible in this image). Expression of mStayGold::GG also enabled visualization of phagocytic cups that form at the base of the collar during bacterial engulfment in asexual cells (asterisk). See also Video S9. (D) In contrast with asexual cells, female gametes have collars (c) that are nearly twice as long and the cell bodies (b) have reduced circularity (Figure S7B-E and Video S13). (E) Male gametes lack a collar, the cell body (b) is reduced in size, and many display a basal projection that we have termed the “fertilopod” (arrow). Scale bar = 5 µm. See also Video S16. (C’) In live asexual cells grown in replete conditions, Cupidon is detected primarily in an ∼1.5 µm subcollar belt (arrowhead; see also F, G) and in the distal ∼1 µm of the collar. See also Video S10. (D’) In female gametes, Cupidon is restricted to a narrow ∼0.5 µm subcollar belt (arrowhead, see also F, G and Video S14) and is detected faintly throughout the collar. (E’) In male gametes, Cupidon is enriched in the fertilopod (arrow). Scale bar = 3 µm. See also Videos S17-18. (F) Representative subcollar belts of Cupidon::mStayGold in an asexual cell (left) and a female gamete (right). Brackets indicate width measurements taken for the analysis in panel G. Scale bars = 2 μm. (G) The subcollar belt of Cupidon::mStayGold is at least four times as wide in asexual cells (N = 213) compared with female gametes (N = 213). Three biological replicates of asexual cells and three of female gametes were analyzed. P-values were calculated from an unpaired two-tailed t-test. (H, I, J) Cupidon::mStayGold is trafficked dynamically along microvilli in the collar. (H) By tracking individual microdomains (blue and orange) over time, we observed both anterograde and retrograde trafficking of Cupidon::mStayGold. Scale bars = 2 μm. (also see Videos S11-12). (I) Representative images showing Cupidon::mStayGold localization in the collar of an asexual cell (left) and a female gamete (right). To avoid merging the signals from superimposed microvilli, only half of the collar was projected in these images. The shorter length of some microvilli observed in both images is an artifact caused by the projection of microvilli that are not parallel to the slide. Scale bars = 3 μm. (J) Cupidon::mStayGold is enriched at the distal end of collar microvilli in asexual cells (light grey; N=71), whereas it is more evenly distributed along the proximal-distal axis in female gametes (dark grey; N=77). Cupidon::mStayGold signal intensity (arbitrary units) was measured along the distalmost 4 µm of the collar. Dotted lines indicate the standard deviation across technical replicates (three for asexual cells and three for female gametes). (K) The fertilopod (filled arrowhead) in male gametes is an actin-based structure. Shown is a live male gamete from a Lifeact::mCherry (Lifeact::mCh)^101^ strain imaged by DIC (left), confocal fluorescence microscopy to detect Lifeact::mCh (middle) and merged (right). Scale bars = 2 μm. See also Figure S7P. (L) The fertilopod of the male gamete (arrowhead) attaches to the subcollar belt of the female gamete. Flagellum (open arrowhead). Scale bar = 5 μm. Also see Video S19. (M) Live imaging of paired *S. rosetta* gametes revealed adhesion between male and female gametes at a Cupidon::mStayGold-enriched fertilopod-subcollar interface (arrowhead). Scale bars = 5 μm. See also Figure S7T and Video S20. (N) Schematic summarizing changes in cell morphology, Cupidon post-translational modifications, and Cupidon localization during the transition from replete to spent medium.

We first assessed whether Cupidon receptor processing might change with nutrient status. The GAIN domains of many choanoflagellate aGPCRs, including *S. rosetta* Cupidon (Figure S5A-B), contain a consensus autoproteolytic cleavage sequence (HL/T) together with additional conserved residues that are implicated in regulating dissociation of the extracellular N-terminal fragment (NTF) from the C-terminal fragment (CTF). In many metazoan aGPCRs, ligand binding or mechanical stimuli^22,35,53–56^ trigger shedding of the NTF, exposing an internal tethered agonist peptide, which then activates the receptor (Figure 3A, Figure S5C-D). Western blot analyses confirmed that Cupidon undergoes cleavage in both replete and spent medium (Figure 3B).

However, the CTF variants detected differed between conditions. In replete cells, we detected a fragment near the predicted size of the cleaved CTF (∼93 kDa; CTF) together with a slightly larger species (∼110 kDa; CTF*), whereas starved cells displayed a distinct higher-molecular-weight fragment (∼115 kDa; CTF**). Because both CTF* and CTF** migrate above the theoretical CTF size, they may correspond to different glycosylated forms of the CTF. In metazoan aGPCRs, glycosylation is a common post-translational modification that controls the trafficking, cleavage, and signaling^57–59;^ Cupidon contains predicted glycosylation sites in both the N-and C-terminal fragments (Figure S6). The full-length receptor (∼220 kDa; FL) was only detected in starved cells (Figure 3B). These data are consistent with nutrient status controlling Cupidon cleavage state and posttranslational modifications, potentially altering receptor activity.

Next, we analyzed the subcellular localization and dynamics of Cupidon::mStayGold in live asexual cells and gametes. In cells grown in replete conditions, the receptor was enriched in an ∼1.5 µm subcollar belt and in the distal ∼1 µm of the collar (Figure 3C, C’, F, G and S7F-G; Videos S9-10). Between these two reservoirs, dynamic microdomains of Cupidon::mStayGold were transported along the microvillar membrane, either towards the tips (anterograde transport; ∼2.3µm/s) or towards the microvillar base (retrograde transport; ∼1.7µm/s) (Figure 3H, S7N-O; Videos S11 and 12). While this localization pattern of Cupidon was also observed in *S. rosetta* chains, rosettes, and thecate cells (Figure S7H-K), it differed in gametes, which have distinct cell architectures (Figure S7L).

In female gametes, the cell body and cell collar are elongated by 61% and 55% relative to asexual cells (Figure S7A-E; Video S13). In these cells, Cupidon was restricted to a ∼0.5 µm subcollar belt (Figure 3D, D’, F, and G; Video S14) and detected in faint microdomains along the collar microvilli, where it was more evenly distributed than in asexual cells but shared similar trafficking dynamics (Figure 3H, I, and J, Figure S7N-O; Videos S14-15). Male gametes lack a cell collar and have a smaller and rounder cell body (Figure S7A-E; Video S16). Here, Cupidon was specifically enriched in a previously undescribed basal projection, hereafter termed the “fertilopod” (Figure 3E, E’, K and S7L-M; Videos S16-18). The fertilopod extends from the basal pole in differentiated male gametes and contains actin filaments. Importantly, male gametes were predominantly paired with female gametes through the fertilopod, which attached to the subcollar belt of female gametes (Figure 3L, S7Q-T; Video S19). Observation of live paired gametes expressing Cupidon::mStayGold confirmed that male and female gametes specifically adhered to each other through these two Cupidon-enriched compartments (Figure 3M, S7T; Videos S20-21).

Together, our results revealed that the transition from replete to spent medium induces changes in cell morphology, Cupidon post-translational modifications, and Cupidon subcellular localization (Figure 3N). These findings establish Cupidon as a nutrient-dependent regulator of collar adhesion. Moreover, localization of Cupidon to the cell collar in asexual cells coincides with the ectopic collar adhesion observed in *cupidon*^1–95^ cells, leading us to hypothesize that Cupidon inhibits adhesive molecules present on the collar of asexual cells. Finally, the enrichment of Cupidon at the contact interface between male and female gametes was counterintuitive, given its role as an inhibitor of cell-cell adhesion under nutrient-replete conditions. This raises the possibility that Cupidon may be repurposed to mediate gamete recognition or fusion under starvation conditions.

### Gamete fusion is regulated by Cupidon

Because loss of *cupidon* under nutrient-replete conditions triggers starvation-like clumping, and because Cupidon protein localizes to the site of contact between gametes, we next investigated whether Cupidon influences mating. To this end, we developed a flow cytometry–based assay for cell fusion. We used four transgenic lines: *cupidon*^wt^ and *cupidon*^1–95^ cells expressing either a cytoplasmic mTFP marker (mTFP-cyto) or a nuclear histone H2B fused to mCherry^60^ (H2B::mCh; Figure S8A). To measure cell fusion, we mixed differentially labeled cells of the same genotype in equal proportions—mTFP-cyto with H2B::mCh—and monitored the frequency of fusion, represented by the percentage of double-positive cells detected by flow cytometry (Figure S8B-D; an example of ongoing cell fusion is shown in Video S22). The experiment was conducted under both replete and starvation conditions. Because this approach only detects fusions between cells carrying different markers, it captures approximately half of all fusion events occurring in a population; reported percentages therefore represent a lower bound of total fusion frequency.

Under replete media conditions, fusion events were undetectable in both *cupidon*^wt^ and *cupidon*^1–95^ (Figure S8E and F), indicating that loss of Cupidon does not promote mating in nutrient-replete conditions. Under starvation conditions, fusion of *cupidon*^wt^ cells increased over time, reaching approximately 3% of the total cell population by Day 5. In contrast, *cupidon*^1–95^ cells showed markedly impaired fusion, with double-positive cells remaining below 0.5% at all time points—a roughly 7-fold reduction relative to wild type (Figure 4 and Figure S8G). These data, together with the localization of the receptor at the site of contact between gametes (Figure 3M), indicate that Cupidon is a key regulator of starvation-induced gamete fusion in *S. rosetta*. This result was unexpected. Because loss of *cupidon* triggers ectopic aggregation under nutrient-replete conditions, we initially hypothesized that Cupidon might be downregulated or inactivated in gametes. Instead, Cupidon is specifically required for fusion under starvation, suggesting it plays a distinct, cell-state-specific role at the gamete contact interface.

**Figure 4.**
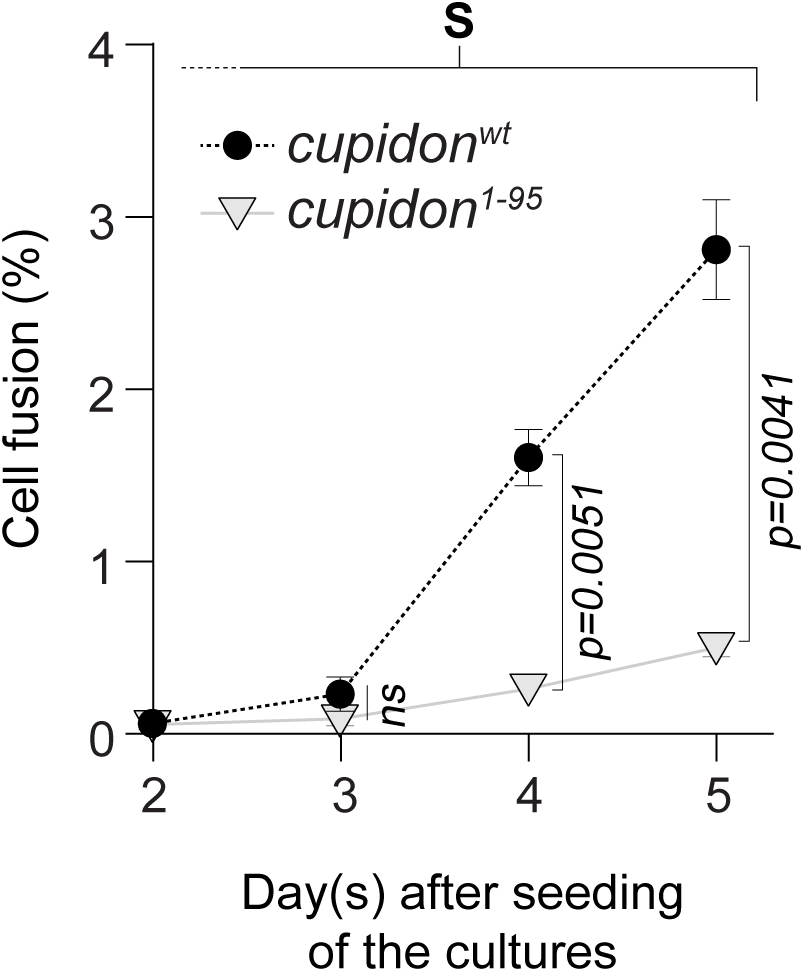
Cupidon is required for proper gamete fusion. In spent medium, cell fusion events in *cupidon^wt^* increased over time, reaching ∼3% of the total cell population by Day 5. In contrast, cell fusion events represented fewer than 0.5% of double-positive cupidon^1–95^ cells by Day 5. Three biological replicates were analyzed per genotype; the mean values were plotted, with standard deviations shown as error bars. A total of 10,000 events were analyzed per replicate for both wild-type and *cupidon*^1–95^ mutant cultures at each time point (see also File S6). P-values were calculated from a two-tailed t-test with Welch’s correction performed independently for each time point.

### GlcNAc-containing glycans mediate collar adhesion and co-localize with Cupidon at the male–female gamete interface

We then investigated possible molecular mechanisms underlying Cupidon’s influence on cell aggregation and mating. Several lines of evidence pointed to a role for surface glycans. Changes in the regulation of surface glycans promote mating and ectopic cell aggregation in *S. rosetta*^20,46^. In addition, the extracellular region of Cupidon contains a predicted sugar-binding ConA-like domain (Figure 1B), suggesting that Cupidon may directly engage cell-surface glycans. Moreover, ALFA-tagged Cupidon cleavage products were detected at unexpectedly high apparent molecular weights on immunoblots (Figure 3B, CTF* and CTF**), consistent with glycosylation, and *in silico* analysis revealed multiple predicted N-and O-glycosylation sites on the receptor (Figure S6).

To assess whether surface glycans could underlie the Cupidon phenotypes, we first examined the glycocalyx of *cupidon*^wt^ and *cupidon*^1–95^ cells under both replete and starvation conditions. We screened a panel of seven fluorescently labeled lectins^46^ (Table S2); all seven lectins produced identical staining patterns in both *cupidon*^wt^ and *cupidon*^1–95^ cells (Figure S9D-E), indicating that loss of Cupidon does not cause global changes in the cell-surface glycan landscape of asexual cells or gametes. Among the lectins tested, two—LEL (*Lycopersicon esculentum* lectin) and STL (*Solanum tuberosum* lectin), which recognize N-acetylglucosamine (GlcNAc) within poly-N-acetyllactosamine (pLacNAc) chains^61^ —showed overlapping distributions with Cupidon::mStayGold (Figure 5A-C, S9A-B). In asexual cells grown in replete conditions, LEL and STL co-localized with Cupidon at the microvillar tips and the subcollar belt; an additional ring of GlcNAc staining was also observed at the microvillar base, where Cupidon staining is not detected (Figure 5A–C, Video S23). In contrast, Jacalin—a lectin that binds galactosyl N-acetylgalactosamine (GalNAc)^61^—showed an anti-correlated localization pattern: Jacalin staining was enriched at the microvillar base, in a broad lateral band below the Cupidon-rich subcollar domain, and in the basal cap, occupying domains distinct from those of Cupidon (Figures 5A, D–E; S9C and Video S24 and S25).

**Figure 5.**
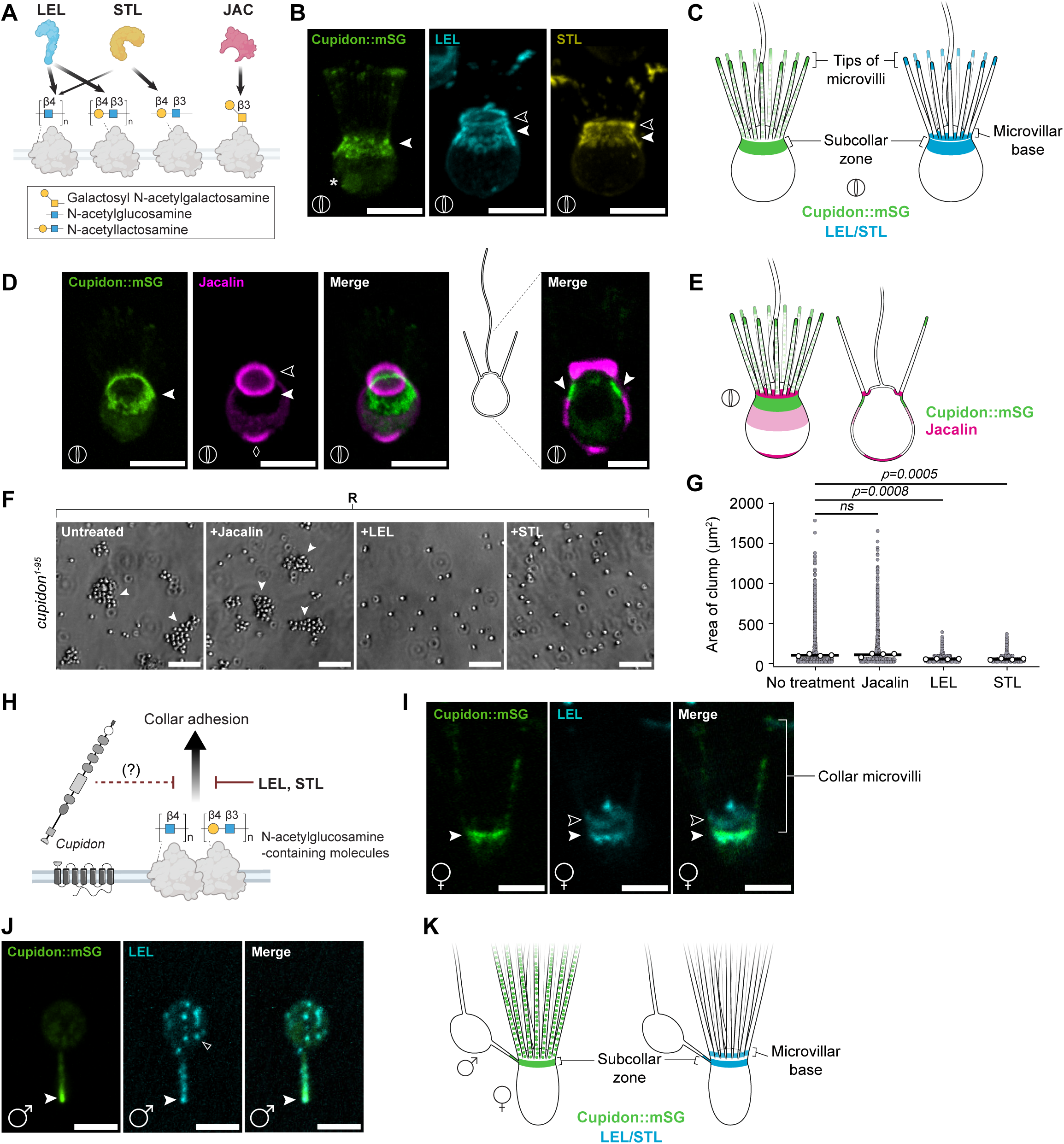
N-acetylglucosamine-containing glycans colocalize with Cupidon and are required for collar adhesion. (A) Schematic illustrating the sugar-binding specificities of the commercially available fluorescently labeled lectins used in this study. LEL (*Lycopersicon esculentum* lectin) and STL (*Solanum tuberosum* lectin) recognize N-acetylglucosamine within poly-N-acetyllactosamine (pLacNAc) chains. Jacalin recognizes galactosyl N-acetylgalactosamine^61^. See also Table S2. (B) In asexual cells, Cupidon::mStayGold (left; green) localizes to a subcollar belt (arrowhead) and to the tips of the collar microvilli. Asterisk marks autofluorescence from the food vacuole. LEL (middle; cyan) and STL (right; yellow) localize to the subcollar belt (arrowhead), the microvillar base (open arrowhead), and at the tip of the microvilli (see Figure S9A-B and Video S23). Scale bars = 5 μm. (C) Schematic summarizing the overlapping distributions of Cupidon::mStayGold (green) and LEL/STL staining (cyan) at the microvillar tips and in the subcollar belt of asexual cells. (D) Cupidon::mStayGold (green) and Jacalin (magenta) occupy non-overlapping domains along the apical-basal axis of asexual cells. Cupidon::mStayGold is enriched in the collar and in the subcollar belt (arrowheads), while Jacalin staining is enriched at the microvillar base (open arrowhead), the lateral domain (bracket), and the basal cap (diamond). The rightmost panel shows the merge in cross-section. Scale bars = 5 µm for projections and 3 µm for cross-section of merge. See also Videos S24-25. (E) Schematic illustrating the complementary distributions of Cupidon::mStayGold (green) and Jacalin (magenta) in asexual cells. (F) Cell clumps formed by *cupidon*^1–95^ cells under replete conditions disperse after treatment with LEL or STL, but not with Jacalin. Arrowheads indicate cell clumps. Scale bars = 50 µm. See also Video S26. (G) Quantification of clump area (µm²) for *cupidon*^1–95^ cells treated as in (F). LEL and STL treatment markedly reduce clump size relative to untreated cells or Jacalin-treated cells. Three biological replicates were analyzed per condition. A minimum of 5374 projected areas (single cells or clumps) were plotted for each condition; white dots represent means per replicate; bars denote means across replicates. P-values were calculated from a one-way ANOVA with Dunnett’s multiple comparisons test. (H) Model of glycan-mediated collar adhesion. N-acetylglucosamine-containing molecules at the cell surface promote collar adhesion. Cupidon suppresses this adhesion through an as-yet-uncharacterized mechanism. Exogenous LEL and STL compete for N-acetylglucosamine-binding sites and thereby inhibit collar adhesion. (I) In female gametes, Cupidon::mStayGold (green) colocalizes with the more basal LEL-positive subcollar belt (arrowhead) but not with the more apical microvillar base (open arrowhead). Scale bar = 3 µm. See also Video S27. (J) In male gametes, LEL staining colocalizes with Cupidon::mStayGold in the fertilopod (arrowhead); puncta of LEL signal are also observed in the cell body (open arrowhead, left). Scale bars = 3 µm. See also Figure S9G-H and Video S28. (K) Model of gamete pairing. During mating, the fertilopod of the male gamete contacts the Cupidon::mStayGold-positive, LEL/STL-reactive subcollar belt of the female gamete.

Because LEL and STL co-localized with Cupidon, we next tested whether GlcNAc-containing molecules are required for collar adhesion in *cupidon*^1–95^ cells. Treatment of *cupidon*^1–95^ cultures with LEL or STL inhibited clump formation under replete conditions, significantly reducing clump area relative to untreated controls (Figs. 5F–G; Video S26). LEL and STL likely act by competitively blocking the GlcNAc-binding sites that mediate cell-cell contact. Jacalin treatment, on the other hand, had no effect on clumping, indicating that the inhibitory activity was specific to GlcNAc-binding lectins rather than a general consequence of lectin addition (Figures 5F–G and Video S26). These results indicate that GlcNAc-containing molecules at the cell surface likely constitute the adhesive substrate driving collar-mediated aggregation in *cupidon*^1–95^ cells. We propose that Cupidon normally suppresses the accessibility or adhesive activity of these molecules under replete conditions (Figure 5H).

Finally, we examined whether the spatial relationship between GlcNAc and Cupidon localization observed in asexual cells is preserved at the gamete contact interface. In female gametes, LEL staining co-localized with Cupidon::mStayGold in the subcollar belt, while also revealing a more apical ring of LEL signal in the microvillar base (Figure 5I, S9F and Video S27). In male gametes, LEL staining was punctate in the cell body and enriched at the fertilopod tip, where it overlapped with Cupidon::mStayGold, although the intensity of this enrichment varied across cells (Figure 5J, S9G-H, and Video S28). As in asexual cells, Jacalin-and Cupidon::mStayGold-enriched territories were distinct in gametes (Figure S9I and Videos S29 and S30). Taken together, these observations indicate that the male–female contact interface is enriched in both Cupidon (Figure 3M, S7L-M and T) and GlcNAc-containing molecules (Figure 5I-J, S9J), consistent with a model in which GlcNAc-mediated adhesion at the fertilopod–subcollar belt junction promotes gamete fusion (Figure 5K).

## DISCUSSION

Here, we identify Cupidon as a regulator of two key steps of mating in *S. rosetta*: cell aggregation and gamete fusion (Figures 1D-E and 4). In nutrient-replete cells, Cupidon suppresses ectopic cell-cell adhesion by inhibiting GlcNAc-dependent collar adhesion (Figure 5F, G). Under starvation, changes in Cupidon processing and localization allow cell aggregation (Figure 2 and Figure 3B-J). Through its enrichment at the contact site between male and female gametes, Cupidon also promotes gamete fusion (Figure 4). Cupidon therefore appears to reprogram the adhesive landscape of the cell surface according to cell state, coupling environmental conditions to the process of mating.

Gametogenesis involves both the relocalization of Cupidon and morphological changes in the gametes. In this study, we uncovered new morphological features of male and female gametes that differ from asexual cells and from each other (Figure 3C-E and Figure S7A-E; Videos S9, S13, and S16). Each possesses Cupidon-enriched gamete-specific structures: the elongated female collar and the male fertilopod (Figure 3C’-E’ and 3F-J; Figure S7G and L; Video S14 and S17). Together, these structures represent a specialized contact interface for gamete pairing.

The collar, normally associated with feeding^62^, elongates and likely contributes to the recruitment of male gametes. The fertilopod adds a second layer of specialization to the gamete interface. Like the sperm acrosomal process in metazoans^63–67^, it is an F-actin-rich protrusion implicated in gamete adhesion and fusion (Figure 3K and 3M, Figure 5, Figures S7 P-T). Whether this structural and functional parallel reflects homology or convergent evolution is an open question, although it raises the possibility of an ancestral role for F-actin-based protrusions in gamete recognition and fusion.

The gamete-specific remodeling of the Cupidon membrane territories described above (Figure 3C’, D’, and E’ and Figure S7G-M) occurs in parallel with a re-organization of cell-surface sugars. In asexual cells, the receptor is largely confined to the subcollar belt and the tips of the collar microvilli, with dynamic Cupidon-positive microdomains moving bidirectionally between these regions (Figure 3C’, F, H, and Videos S10-12). The overlap between Cupidon-rich regions and GlcNAc-containing molecules, together with the anticorrelation of these regions with GalNAc-containing molecules (Figure 5B, D), suggests that the collar membrane is partitioned into different glycan-defined territories. This partitioning may constrain Cupidon localization and activity.

This regional organization shapes how and where cells adhere across different physiological states. In nutrient-replete *cupidon* mutants, ectopic adhesion occurs primarily through the collar, the same domain where Cupidon normally suppresses GlcNAc-dependent adhesion (Figures 2E and F). In starved wild-type female gametes, collar-mediated adhesion is similarly predominant, consistent with the hypothesis that changes in Cupidon processing relieve this suppression. By contrast, EroS-treated wild-type cells adhere primarily through the cell body, suggesting that this pathway induces a distinct adhesive mechanism operating independently of the collar. In sexual pairing, the male fertilopod engages the female subcollar region in a stereotyped manner (Figure 3L, M), adding a third mode of contact that is spatially precise and cell-state specific. Thus, adhesion in *S. rosetta* is a regionally specialized and cell-state-dependent property of the cell surface. Cupidon is perhaps best understood as a regulator of specific adhesive territories: its redistribution and altered processing during cell-state transitions help determine which cell-surface interfaces are adhesive and when.

GlcNAc may provide the molecular link between Cupidon’s anti-adhesive role in asexual cells and its pro-mating role in sexual cells. GlcNAc is well known as a structural component of cell surfaces and extracellular matrices^68,69^, but it also contributes to sperm-egg recognition in diverse metazoans^70–76^. The enrichment of GlcNAc-containing molecules at the *S. rosetta* gamete contact interface (Figure 5I, J; Figure S9F-H, J) suggests that glycan-based recognition may play a similar role here. The model that Cupidon represses GlcNAc-dependent adhesion in asexual cells is supported by two complementary lines of evidence: the co-localization of GlcNAc-binding lectins (LEL and STL) with Cupidon at the collar (Figure 5), and the competitive inhibition of cell clumping by the same lectins, consistent with blocking GlcNAc-mediated cell contact (Figure 5F, G). Together, these findings suggest that Cupidon regulates a glycan-based adhesive system whose output depends on developmental context.

To our knowledge, Cupidon is the first aGPCR to be functionally characterized outside metazoans. Cupidon shares defining architectural features of metazoan aGPCRs, including a large adhesion-rich extracellular region, a cleavable GAIN domain, a canonical 7TM domain, and a predicted PDZ-binding motif (Figure 1B). Like metazoan aGPCRs, it undergoes regulated proteolytic processing and spatially patterned membrane localization^32,77,78^ (Figure 3B-E). Together, these observations suggest that aGPCR-mediated adhesion predates metazoan multicellularity and that *S. rosetta* uses this ancient receptor architecture to regulate sexual reproduction.

Two questions raised by this work remain open. First, the basis of the fusion defect in *cupidon*^1–95^ gametes remains unresolved. Because the high motility of gametes precluded long-term imaging of individual pairing events, we cannot currently determine whether the reduced fusion frequency reflects a direct role for Cupidon in the fusion machinery or an indirect consequence of impaired gamete pairing; improved microscopy and microfluidic approaches to confine gametes will be needed to resolve this. Second, the extracellular signal that triggers Cupidon processing and relocalization during starvation is unknown. Like many metazoan aGPCRs, Cupidon may be activated through the exposure of an otherwise hidden tethered agonist following the shedding of its N-terminal region (Figure S5), but whether the initiating cue is a nutrient, a bacterially derived molecule, a stress signal, or a contact-dependent ligand presented by a partner cell remains to be identified.

Cupidon illustrates how a single receptor can regulate multiple steps of a developmental process by coupling environmental changes to the spatial remodeling of the cell surface. More broadly, the molecular parallels between Cupidon-mediated mating in *S. rosetta* and fertilization in metazoans suggest that the molecular logic underlying gamete recognition and fusion may predate the origin of metazoans.

## MATERIAL AND METHODS

### *S. rosetta* culture conditions

All experiments were performed using wild-type (SrEpac) or transgenic strains of *S. rosetta* co-cultured with *Echinicola pacifica*^50^. Throughout this study, cells were grown either in <u>n</u>utrient-replete (NR) conditions, which promote the growth of asexual cells, or in <u>s</u>pent media/<u>s</u>tarvation conditions (S), which trigger the sexual cycle and the production of male and female gametes. Although these methods result in distinct outcomes, both use the same initial culture medium, (hereafter referred to as “1.5%/1.5% medium,”) but differ in culturing duration and in the density of *E. pacifica* prey bacteria provided in the cultures, as described below.

The standard 1.5%/1.5% culture medium, previously described as low nutrient media (LNM)^15^, consists of AK sea water (AKSW) complete (AKSWC; 5.0 g/L peptone, 3.0 g/L yeast extract, and 3.78 g/L glycerol dissolved in AK seawater) and AK cereal grass media (AKCGM3; 5 g/L cereal grass steeped in AKSW (Carolina Biological Supply Company, Cat. No. 132375). Both were diluted to 1.5% by volume in AKSW and supplemented with nitrates, phosphates, trace elements, and vitamins as specified in Booth and King (2020)^15^. This medium was preferred as it prevents *E. pacifica* biofilm formation during routine passaging while supporting cell densities sufficient for experimental use.

To grow asexual cells, cultures were passaged every two days at a 1:500 dilution into fresh 1.5%/1.5% culture medium supplemented with 90 µg/mL *E. pacifica* prey bacteria (from a 10 mg/mL *E. pacifica* suspension in ASW) to ensure a consistent supply of prey bacteria. Cultures were maintained at 22°C and 60% humidity. One day prior to experiments, a mid-log phase culture (between 1 × 10^5^ and 1 × 10^6^ cells/mL), passaged 48 hours earlier, was diluted 1:30 into 1.5%/1.5% culture medium containing 150 µg/mL *E. pacifica*. After 24 hours of incubation at 22°C and 60% humidity, the cells were pelleted by centrifugation (2000 × g for 10 minutes at 22°C), resuspended in ∼1 mL residual volume, and the cell density was measured. These cells were then diluted to 1 x 10^5^ cells/mL in AKSW and used immediately.

To promote starvation-induced mating and gamete production, cells were maintained for 3 to 4 days without passaging. Under these conditions, the *E. pacifica* prey bacteria was progressively depleted by the choanoflagellates and the bacterial density dropped dramatically, coinciding with the production of male and female gametes. On the day of the experiment, the culture was pelleted by centrifugation (2400 × g for 10 minutes at 22°C), resuspended in the residual volume (∼1 mL), and the cell density was measured. The cells were then diluted to 1 x 10^5^ cells/mL in AKSW and used immediately.

Preparation of cell cultures for RNAseq analysis and cell fusion experiments differed slightly from the protocols described above and is explained in the “RNAseq analysis” and “Cell fusion assay” sections, respectively.

### Growth curve assay

One day before the start of the growth assay, log-phase cultures were diluted 1:30 into 1.5%/1.5% medium supplemented with 100 µg/mL *E. pacifica* to standardize growth conditions across starter cultures. On the day the growth curve plates were seeded, starter cultures were washed three times with ASW to remove bacteria on a Thermo Fisher Scientific ST16R swinging-bucket rotor centrifuge (2,400 × g for 5 min each), thereby standardizing nutrient conditions across cultures. Cells were then diluted to 5,000 cells/mL in 1.5%/1.5% culture medium and supplemented with 267 µg/mL *E. pacifica* pellet^15^. 500 μL aliquots of culture were transferred into each well of a 24-well plate (Fisher Scientific, Cat. No. 09-761-146) and incubated at 22°C and 60% humidity. Plates were maintained in a covered plastic container with dampened paper towels and a loosely affixed lid to minimize evaporation while allowing gas exchange.

Every 12 hours for 96 hours, three wells from each strain were fixed with 10 μL of 16% paraformaldehyde (Fisher Scientific Cat. No. 50-980-487) and stored at 4°C. After all time points were collected, each sample was vortexed at high speed for 15 seconds to ensure full mixing, and 10 μL was transferred onto a counting slide (Logos Biosystems Cat. No. L12001). The cells were counted using a Luna-FL automated cell counter (Logos Biosystems, Cat. No. L20001; Figure 1C).

### Quantification of cell clump formation

We adapted a protocol developed by Woznica *et al*. (2017)^20^ and Wetzel *et al*. (2018)^46^ to quantify clump size. Cells were diluted to 5 × 10 ^^5^ cells/mL in AKSW and vortexed for 15 s to disrupt preformed clumps. Aliquots of 300 µL were plated in a chambered coverslip (µ-Slide 8 Well ibiTreat, Cat.No. 80826) and incubated for 30-60 minutes under a Zeiss Axio Observer.Z1/7 Widefield microscope equipped with a Hamamatsu Orca-Flash 4.0 LT CMOS Digital Camera (Hamamatsu Photonics) at room temperature (∼22°C).

For measuring clumps (Figure 2F-G, 5F-G, S2E-F) or rosettes (Figure 1F-H), the cultures were imaged at 10 distinct locations throughout the well using a Plan-Apochromat 10x/0.45 Ph 1 M27 objective with a 1.6x Optovar setting.

For time-lapse imaging of clump formation in *cupidon^wt^* and *cupidon* mutant cultures (Figure 1D-E), the cells were allowed to settle for 5 minutes in uncoated wells before being imaged every 30s during 3 hours with a Plan-Apochromat 10x/0.45 Ph 1 M27 objective with a 1.6x Optovar setting at 4 distinct positions per well.

The resulting images were batch-processed in Fiji^79^ for consistency. To reduce background bacterial signal, the Process>*Smooth* command was applied, followed by *Process* > *Find Edges* to detect the phase-bright boundaries of the choanoflagellate cells. Images were converted to binary using the *Process>Binary> Make Binary* command, followed by the *Process>Binary>Close-* command to fill in small holes. Finally, images were analyzed using the *Analyze > Analyze particles* command to calculate the area of the binary masks; only particles larger than 20 μm^2^ were kept, filtering out any remaining bacterial signal. Data were plotted in GraphPad Prism (v. 11.0.0) as scatter plots, showing the mean (horizontal line) and standard deviation.

### Western blot analysis of Cupidon post-translational processing

To evaluate the post-translational regulation of Cupidon (Figure 3B), cultures were seeded at 8,000 cells/mL in 120 mL of 1.5%/1.5% medium supplemented with 167 µg/mL *E. pacifica* prey bacteria (from a stock at 10 mg/mL *E. pacifica* suspension in ASW) in 3-layer culture flasks (Falcon™ Cell Culture Multi-Flask, 3-layer, 525 cm^2^, Corning, Cat. No. 89204-476) kept at 22°C and 60% humidity. Cells were collected after 38 hours (for asexual cells) or 82 hours (for gametes).

The cultures were centrifuged and washed three times with AKSW (by centrifuging at 2000 × g for 10 minutes for the first wash, 2100 × g for 5 minutes for the second, and 2300 × g for 5 minutes for the third) to minimize bacterial contamination. Aliquots of 1 x 10^7^ cells were pelleted and gently resuspended in non-selective lysis buffer (1X DTT (1mM final), 1X cOmplete^™^, Mini, EDTA-free Protease Inhibitor Cocktail (Roche Cat. No. 11836170001; from a 20 X stock diluted in 1.12X Triton Lysis buffer), 1X Benzonase Nuclease (Millipore Cat. No. 70746; from a 450 X stock at 25-29 U/µL), 5% (v/v) of Milli-Q water, and 89% (v/v) of 1.12X Triton Lysis buffer (56mM Tris pH 8, 168mM NaCl, 1.12mM EDTA, 0.112mM EGTA, ∼2.24% Triton X-100). Lysis proceeded for 20 minutes on ice with vortexing for 1 second every 5 minutes. The cells were then centrifuged at 18,000 × g for 15 minutes at 4°C, and the collected supernatants were either used directly or flash-frozen and stored at-80°C for later use.

Protein concentration was determined by Bradford assay (Thermo Fisher, Cat. No. 23200) using a NanoDrop 2000c. Samples (20 µg of total protein) were denatured in 4X Protein Loading Buffer (LICORbio, P/N: 928-40004) and a final concentration of 100 mM 2-β-mercaptoethanol (Sigma, Cat. No. M3148) in a total volume of 40 µL. The samples were incubated at 95°C for 5 minutes before loading the totality of the volume (40 µL) into a Novex™ Tris-Glycine Mini Protein Gel, (8%, 1.0 mm, WedgeWell™ format; Invitrogen, Cat. No. XP00080BOX) with 10 µL of PageRuler™ Plus Prestained Protein Ladder (Thermo Fisher Scientific, Cat. No. 26619). Gels were run on a Mini Gel Tank (Thermo Fisher Scientific, Cat. No. A25977) in 1X Novex™ Tris-Glycine SDS Running Buffer (Thermo Fisher, Cat. No. LC2675) at 225 V for 40 minutes at room temperature. The gels were then transferred to a 0.45 µm nitrocellulose membrane (Bio-Rad Cat. No. 1620115) in transfer buffer (1X Tris-Glycine Transfer buffer; Thermo Fisher Scientific, Cat. No. LC3675, with 0.05% SDS and 10% Methanol) for 20 hours at 4°C (10 V, 70 mA) using Mini Blot Modules (Thermo Fisher Scientific, Cat. No. B1000).

The membranes were then briefly washed in ultra-pure water for 1 minute, air-dried for 1 to 2 hours at room temperature to improve image quality, rehydrated in 1X TBS (Thermo Fisher Scientific, Cat. No. J60764.K2) for 5 minutes on an orbital shaker and blocked for 1 hour at room temperature with Intercept (TBS) Blocking Buffer (LICORbio, P/N: 927-60001). The membranes were then incubated for 20 hours at 4°C with anti-Tubulin [YOL1/34] (abcam, Cat. No. ab6161) diluted to 1 µg/mL in Intercept T20 (TBS) Antibody Diluent (LICORbio, P/N: 927-65001). The membranes were then washed once in TBS-T (TBS with 0.1% Tween 20) by orbital shaking at room temperature for 5 minutes and subsequently immunostained with FluoTag-X2 anti-ALFA LI-COR IRDye 800CW (NanoTag Biotechnologies Cat. No. N1502-Li800-L) diluted 1:10,000 and IRDye® 680RD Goat anti-Rat IgG Secondary Antibody (LICORbio, P/N: 926-68076) diluted 1:5,000 in Intercept T20 (TBS) Antibody Diluent. The membranes were incubated for one hour at room temperature on an orbital shaker before washing them three times (5 minutes each) with TBS-T. A final wash of the membranes in 1X TBS for 5 minutes was performed before imaging them on a Li-Cor Odyssey CLx Imaging System in the 700nm and 800nm channels with the following settings: Setup: Membrane, Channels: Auto (700 and 800), Scan Controls: 84µm or less. This Western blot protocol has been adapted from a previous study, Rutaganira *et al*. (2025)^80^, and is also described in part in Leon et al (2025)^81^.

### Cell state transitions and cell-cell interactions

#### Rosette induction with RIF-OMVs

To induce rosette formation in cultures, we used RIF-OMVs from the bacterium *Algoriphagus machipongonensis* (ATCC, BAA-2233) as previously described^18,82^. Briefly, the cultures were prepared at 1 x 10^4^ cells/mL into a final volume of 3 mL of 1.5%/1.5% culture medium supplemented with 60µg/mL *E. pacifica* food bacteria and incubated either with RIF-OMVs at a dilution of 1:4000 or with an equal volume of 50mM HEPES-KOH pH 7.4, the dilution buffer used to resuspend RIF-OMVs. Each culture was then transferred to a 6-well plate (CELLSTAR® 6 Well Cell Culture Multiwell Plates, Item No.: 657185) and incubated for 48 hours at 22°C and 60% humidity.

After a 48-hour incubation period, the live cultures (Figure S2D and Video S4) were imaged by transferring 200 μl of cells into a chambered coverslip (µ-Slide 8Well ibiTreat, Cat.No. 80826) using a wide-bore pipette tip for minimal disruption, and allowing them to settle for 15 min. For images of vortexed RIF-OMVs-induced and uninduced cultures (Figure 1G-H), 200µl aliquots of cells were fixed for 10 minutes at room temperature with 50µl of 16% paraformaldehyde (Fisher Scientific Cat. No. 50-980-487), then vortexed vigorously for 15 s before transferring into a chambered coverslip (µ-Slide 8Well ibiTreat, Cat. No. 80826) coated with 0.1 mg/mL poly-D-lysine (Millipore Sigma, Cat. No. P6407) for 5 min and washed 3 times with ASW to remove excess poly-D-lysine. The fixed cells were allowed to settle for 10 minutes before imaging. Fixed and live cells were imaged under a Zeiss Axio Observer.Z1/7 Widefield microscope with a Hamamatsu Orca-Flash 4.0 LT CMOS Digital Camera (Hamamatsu Photonics) and a Plan-Apochromat 10x/0.45 Ph 1 M27 objective with a 1.6x Optovar setting (Figure 1G and Figure S2D). The quantification of rosette induction and their resistance to shear stress (Figure 1G-H) was performed on images of fixed, vortexed cells using the protocol described under the section “Quantification of cell clump formation”.

#### EroS_pv_-induced mating clumps

Cell cultures diluted to 5 × 10^5^ cells/mL in AKSW were treated with 0.0035 units of EroS_pv_^20^ (Chondroitinase ABC from *Proteus vulgaris*, Sigma-Aldrich, SKU: C3667) for 15 min (Figure 2B-E and Figure S3D) or 1 hour (Figure 2F-G) to induce mating clump formation at room temperature (∼22°C) before imaging. The quantification of clumps was performed using the protocol described under the section “Quantification of cell clump formation”.

#### Collar depolymerization with Latrunculin A

To assess the effect of the loss of collar on cell clumping, cultures were prepared at 5 x 10^5^ cells/mL into a final volume of 200µl in AKSW supplemented with 5 µM of Latrunculin A (Thermo Fisher Scientific, Cat. Number: L12370) or an equivalent dilution of DMSO, the solvent used to resuspend Latrunculin A. The cultures were then vortexed vigorously for 15 s before being plated onto a chambered coverslip (µ-Slide 8Well ibiTreat, Cat.No. 80826) and incubated directly under a Zeiss Axio Observer.Z1/7 Widefield microscope equipped with a Hamamatsu Orca-Flash 4.0 LT CMOS Digital Camera (Hamamatsu Photonics) for 1 hour at room temperature (∼22°C). After a 1-hour incubation, during which the clumps were allowed to form without mechanical disturbance, the live cells were imaged with a Plan-Apochromat 20x/0.80 Ph 2 M27 objective with a 2.5x Optovar setting (Figure 2F) or a Plan-Apochromat 10x/0.45 Ph 1 M27 objective with a 1.6x Optovar setting (Figure 2G). Clumping was then assessed as described under the section “Quantification of cell clump formation”.

### Lectin-mediated inhibition of cell clumping

Cultures of asexual *cupidon^1-^*^95^ mutant cells were prepared at 5 x 10^5^ cells/mL in AKSW and supplemented with or without Jacalin-Fluorescein (Vector Laboratories, SKU: FL-1151-5), *Lycopersicon esculentum* (Tomato) Lectin (LEL, TL)-DyLight 594 (Vector Laboratories, SKU: DL-1177-1), or *Solanum tuberosum* (Potato) Lectin (STL, PL) (Vector Laboratories, SKU: L-1160-5) at a working concentration of 100ng/mL. The cultures were then vortexed vigorously for 15 s, and 200 µL from each condition was transferred into a Corning® 96-well Clear Flat Bottom Ultra-Low Attachment Microplate (Product Number 3474), specifically designed to minimize protein adsorption, which otherwise immobilizes the lectin-coated cells to the bottom of the plate and precludes this assay. The cells were incubated for 30 min at room temperature (∼22°C) under a Zeiss Axio Observer.Z1/7 Widefield microscope equipped with a Hamamatsu Orca-Flash 4.0 LT CMOS Digital Camera (Hamamatsu Photonics). After 30 minutes, during which the clumps were allowed to form without mechanical disturbance, the cultures were imaged with a Plan Apochromat 10x/0.45 Ph 1 M27 objective with a 1.6x Optovar (Figure 5F-G, Video S26). Clumping was then assessed as described under the section “Quantification of cell clump formation”.

### Classification of cell-cell contact types

The cultures were prepared at 5 x 10^5^ cells/mL in AKSW, vigorously vortexed for 15 s, and 200 μl of each culture was seeded into a chambered coverslip (µ-Slide 8Well ibiTreat, Cat.No. 80826) that we directly incubated for 10 minutes at room temperature (∼22°C) under a Zeiss Axio Observer.Z1/7 Widefield microscope equipped with a Hamamatsu Orca-Flash 4.0 LT CMOS Digital Camera (Hamamatsu Photonics). After 10 minutes, during which the clumps were allowed to form without mechanical disturbance, the cultures were imaged with a 100X/1.40 Plan-Apochromat oil DIC M27 objective with a 1x Optovar. The identity of the cell-cell contact was then manually assessed in Fiji, and the data were plotted using GraphPad Prism 11.0.0.

### Cupidon sequence analysis and structural analysis

#### Identification of Cupidon homologs

Cupidon homologs were searched across the choanoflagellate and eukaryote phylogenies using two complementary approaches. First, the full-length *S. rosetta* Cupidon protein sequence was blasted against the entire EukProt v3 dataset^83^ (993 species); top blast hits with E values < 1 x 10^-10^ and query coverage > 50% were considered for further analysis. The 8 validated choanoflagellate ortholog sequences retrieved from this analysis were then aligned together with the full-length *S. rosetta* Cupidon protein sequence using MAFFT^84^ (Version 7.4) with the E-INS-i algorithm). To search for more distantly related homologs, we used the resulting alignment to build an HMM profile of the full-length protein with the *hmmbuild* module of HMMER^85^ v3.3. The resulting HMM was then searched against both Reference proteomes and UniProt proteomes using the hmmsearch algorithm on the HMMER web server interface^86^ with an E-value of 0.001 (https://www.ebi.ac.uk/Tools/hmmer/search/hmmsearch). The nine Cupidon orthologous protein sequences are provided in File S1.

#### Protein domain identification in Cupidon

To identify protein domains in Cupidon and its orthologs, we first analyzed protein sequences using InterProScan^87^ and CDvist^88^. Additionally, we scanned the protein sequences for conserved protein domain HMM profiles with the *hmmscan* algorithm (running on the Pfam 37.2 HMM database with Gathering threshold) on the HMMER web server interface.

Because these methods of detection rely on previously identified protein domains and could therefore miss undiscovered domains or poorly conserved domains at the sequence level, we systematically tested for the presence of conserved folds in Cupidon proteins using AlphaFold (see section “Structural prediction of Cupidon” below). The protein domain annotations for Cupidon and its orthologs are presented in Figure S1A.

#### Structural prediction of Cupidon

We generated predicted structural models of Cupidon proteins using either AlphaFold2 ColabFold^89^ v1.6.1 (num-relax=1;msa_mode=mmseqs2_uniref_env; model_type=alphafold2_ptm; num_recycles=3) or the AlphaFold3 server^90^ with default parameters (https://alphafoldserver.com/). Because AlphaFold structural prediction of large multidomain proteins can be challenging, with lower accuracy and longer processing times, we ran structural analyses on both the full-length protein and truncated protein fragments. The full extracellular region, the full intracellular region, the 7TM domain, or individual fragments from the extracellular region were therefore submitted independently. Five models were generated per input, of which only the top-ranked prediction (based on the ranking_score metric) was selected for downstream analyses. The quality of the predicted structure was assessed based on the pLDDT confidence score; only models with pLDDT scores above 70 were retained.

To find structural homologs of Cupidon protein domains, we searched the AlphaFold-predicted structures against the AFDB-PROTEOME, AFDB-SWISSPROT, AFDB50, CATH50, MGNIFY_ESM30, and PDB100 databases using the search program Foldseek^91^ (https://search.foldseek.com/search)with default parameters. Structural top hits with an associated E-value<1e^-10^ were considered putative structural homologs. Structures were analyzed, and figures were prepared with ChimeraX^92^.

#### Comparison of Cupidon and metazoan GAIN domains

The C-terminal region of the predicted GAIN domain sequences was extracted from the Cupidon protein and its orthologs, along with those of a variety of metazoan aGPCR GAIN domains spanning a representative range of organisms. All the extracted sequences were then aligned on Geneious Prime v2024.07 using MAFFT with L-INS-I algorithm. The complete set of GAIN domain sequences used is provided in File S5.

To compare the AlphaFold-predicted model of the *S. rosetta* Cupidon GAIN domain with metazoan GAIN domains, we retrieved four experimentally solved structures of metazoan GAIN domains — including human BAI3 (4DLO), human GPR110 (8HC0), mouse GPR56 (5KVM), and zebrafish GPR126 (6V55) — all publicly available on the RCSB PDB database (https://www.rcsb.org/)^93^. All these structures were then color-coded and aligned with ChimeraX using the *matchmaker* command.

#### Conservation and distribution of the Cupidon Mystery Domain

The prediction of a high-confidence fold, named Mystery Domain (MD), in the N-terminal region of Cupidon, required further characterization. To assess whether MD could be conserved outside of choanoflagellates, we conducted both sequence and structural searches using the extracted MD domain as a query. BLASTP^94^ searches against both the NCBI and the EukProt databases, together with Foldseek search, revealed that the MD domain is detected in various eukaryotes (choanoflagellates, metazoans, metamonads, Haptista, and SAR), bacteria, and viruses. Top structural hits (E-value<1e^-10^) were color-coded and aligned with ChimeraX. A list of all MD protein sequences used in these analyses is provided in File S3.

### Prediction of Cupidon glycosylation sites

N-and O-glycosylation sites in Cupidon protein sequence were predicted using a combination of computational tools. NetNGlyc-1.0^95^ (https://services.healthtech.dtu.dk/services/NetNGlyc-1.0/, default threshold 0.5), N-GlycoSite^96^ (https://www.hiv.lanl.gov/content/sequence/GLYCOSITE/glycosite.html, default threshold >60%), and GlycoEP^97^ (https://webs.iiitd.edu.in/raghava/glycoep/index.html, BBP method with default threshold 0.0) were used to predict N-glycosylation sites. NetOGlyc-4.0^98^ (https://services.healthtech.dtu.dk/services/NetOGlyc-4.0/, default threshold 0.5), O-GlcNAcPRED-DL^99^ (https://oglcnac.org/pred_dl/input_fasta, threshold >0.6), GlycoEP (https://webs.iiitd.edu.in/raghava/glycoep/index.html, CPP method with default threshold 0.0) were used to identify putative O-glycosylation sites. To be recognized as a predicted glycosylation site, the glycosylation sequence should be detected by at least two of the three predictors for each class of glycosylation. The resulting glycosylation map of Cupidon is presented in Figure S6.

### Cell fusion assay

#### Nutrient-replete conditions

To assess cell fusion under nutrient-replete conditions (Figure S8E and F), cultures were prepared as follows: five strains — (1) SrEpac (unedited *S. rosetta* wild-type strain grown with *E. pacifica*), (2) *mTFP-cyto, rpl36a^P56Q^*, (3) *H2B::mCherry, rpl36a^P56Q^*, (4) *mTFP-cyto, rpl36a^P56Q^, cupidon^1-^*^95^, and (5) *H2B::mCherry, rpl36a^P56Q^, cupidon*^1–95^ — were individually passaged every two days for one week at a 1:500 dilution into fresh 1.5%/1.5% culture medium supplemented with 90µg/mL *E. pacifica* food bacteria and 80 µg/mL of Puromycin (Thermo Fisher Scientific, Cat. No. A1113803) at 22°C and 60% humidity. This was important to ensure that all five cultures had roughly the same choanoflagellate and bacterial densities at the start and had not recently undergone starvation. Each culture was further visually assessed under an inverted microscope to verify that the cells exhibit the morphology and dynamics of asexual cells. One day before setting up the experiment, each of these mid-log-phase cultures was diluted 1:30 into 1.5%/1.5% culture medium containing 150 µg/mL *E. pacifica* food bacteria. After 24 hours of incubation at 22°C and 60% humidity, on the day of the experiment, each culture was pelleted by centrifugation (2000× g for 10 minutes at 22°C), resuspended in 1 mL residual volume, vortexed vigorously for 15 s, and the cell density was measured.

An aliquot of each culture was diluted to 1 x 10^5^ cells/mL in 0.2 µm syringe-filtered 1.5%/1.5% culture medium and directly analyzed on an Attune CytPix Flow Cytometer (Thermo Fisher Scientific). This step is necessary before combining the cultures to determine the exact volumes of both mTFP-and mCherry-positive cells (in both *cupidon^wt^* and *cupidon*^1–95^ genetic backgrounds) to mix in order to achieve a 1:1 fluorescent ratio detected by the Attune CytPix. Indeed, although the clonal fluorescent strains have stably integrated the mTFP-cyto or H2B::mCherry fluorescent transgenes (see “Transfection” section), the signal intensity can vary from cell to cell in these cultures, possibly reflecting various degrees of gene silencing.

Percentages of weakly fluorescent cells, counted as fluorescently negative by the Attune Cytpix (which depends on both the gating strategy and detector sensitivity used), ranged from 6% to 60% in those cultures (File S6). We used the unedited SrEpac strain as a negative control to adjust the voltage settings for both channel colors: BL1, excited with a 488nm laser and equipped with a 510/10 nm filter to detect mTFP fluorescence, and YL2, excited with a 561nm laser and equipped with a 615/25 nm filter to detect mCherry fluorescence (see File S6). This allowed us to delimit four quadrants, each delineating a distinct population of cells either (1) negative for both mTFP and mCherry fluorescence, (2) positive for mTFP but negative for mCherry fluorescence, (3) negative for mTFP but positive for mCherry fluorescence, and (4) positive for both mTFP and mCherry fluorescence (File S6).

Next, we diluted each of the original cultures (resuspended in 1 mL residual volume) at 400,000 cells/mL in 0.2 µm syringe-filtered 1.5%/1.5% culture medium. The *mTFP-cyto, rpl36a^P56Q^*, and H2B::mCherry, *rpl36a^P56Q^* strains from one side, and the mTFP-cyto, *rpl36a^P56Q^*, *cupidon*^1–95^, and *H2B::mCherry, rpl36a^P56Q^, cupidon*^1–95^ strains from the other, were then mixed at a 1:1 fluorescent ratio in a final volume of 15mL, adjusted under the Attune CytPix Flow Cytometer (see File S6). The two resulting 1:1 cultures (both at 400,000 cells/mL in 15mL) were supplemented with 80 µg/mL of Puromycin and 200 µg/mL of *E. pacifica* food bacteria to ensure the cells would not starve during the assay. We vigorously vortexed the cultures for 15 s and distributed 3 x 2 mL of each culture (*cupidon^wt^* and *cupidon*^1–95^) in a 12-well plate (Fisher Scientific, Cat. No. 07-200-82); 2mL were transferred in each well to allow three replicates per condition (*cupidon^wt^*and *cupidon*^1–95^).

After an incubation period of 20 hours at 22°C and 60% humidity, we verified under an inverted microscope that the cultures were still exclusively composed of asexual cells. Each well was then vigorously pipetted before being diluted to 1 x 10^5^ cells/mL in 3 mL of 0.2 µm syringe-filtered 1.5%/1.5% culture medium and directly analyzed on an Attune CytPix Flow Cytometer. Samples were vortexed at full speed for 15 s just before the analysis to prevent cell adhesion; a total of 10,000 events per condition was assessed using a 25 µl/min flow rate. The resulting density plots were analyzed using FCS Express 7 Professional Standalone Research Edition version 7.30.0018 (De Novo Software) (File S6). The averaged ratios of mTFP-to H2B::mCherry-labeled cells depicted in Figure S8C and S8D were calculated by dividing for each of the three replicates the percentage of mTFP-positive cells by the percentage of mCherry-positive cells. The mean and standard deviation were then plotted using GraphPad Prism 11.0.0.

#### Starvation conditions

A similar protocol to the one described above was used to assess gamete fusion under starvation, with the main difference being that cultures were seeded at a final density of 10,000 cells/mL rather than 400,000 cells/mL on the first day of the experiment. Additionally, the cells were monitored for 5 days rather than just 20 hours after the cultures were set up at 22°C and 60% humidity. Because cell morphologies and their associated autofluorescence levels changed over the course of starvation, shifting progressively from a predominantly homogeneous population of asexual cells to a heterogeneous population of differentiated gametes, accurate negative controls, reflecting the state of the culture after a given period of incubation, were required for each day of the time course to adjust the detection settings for both fluorescent channels. Therefore, along with three 1:1 fluorescent culture replicates for both *cupidon^wt^* and *cupidon*^1–95^ conditions per day, an additional SrEpac culture was prepared for each of the 5 days at the same density as the 1:1 fluorescent cultures (10,000 cells/mL in 2 mL), serving as their corresponding negative control. Samples were acquired and analyzed as explained in the previous section.

### Lectin-based glycocalyx profiling

We used an array of FITC-labeled lectins — including Wheat Germ Agglutinin (WGA) (Vector Laboratories, SKU: FL-1021-5), Concanavalin A (Con A) (Vector Laboratories, SKU: FL-1001-25), Jacalin (Vector Laboratories, SKU: FL-1151-5), *Lycopersicon esculentum* (Tomato) Lectin (LEL, TL) (Vector Laboratories, SKU: FL-1171-1), *Maackia amurensis* Lectin I (MAL I) (Vector Laboratories, SKU: FL-1311-2), *Solanum tuberosum* Lectin (STL/PL) (Vector Laboratories, SKU: FLK-4100), and *Vicia villosa* Lectin (VVL, VVA) (Vector Laboratories, SKU: FL-1231-2)—to map the glycocalyx of *S. rosetta* asexual cells and gametes (see Table S2). These lectins were previously shown to be a valuable tool for monitoring changes in cell-surface glycans in *S. rosetta*^46^. Cultures were mixed with each lectin at a 10 µg/mL final concentration, transferred onto a poly-D-lysine–coated 96 well plate (µ-Plate 96 Well Square ibiTreat, Ibidi, ref. 89626), and centrifuged at 200 × g for 1 minute at room temperature using a swinging-bucket rotor to allow the cells to gently adhere. After the centrifugation, the cells were imaged immediately using a Nikon Ti2 spinning-disk confocal system equipped with a Hamamatsu ORCA-Fusion Digital CMOS C14440-20UP camera. Z-stacks (0.2 µm intervals) were acquired using a Plan Apo λ 100× (NA 1.45) MRD01905 oil immersion objective.

### Quantification of cell and collar morphology

To measure the area, aspect ratio, and circularity of the cell body, as well as the length of the collar microvilli (Figure S7A-E), we prepared cultures of mStayGold::GG-expressing asexual cells or gametes at 5 x 10^5^ cells/mL into a final volume of 200 µL in AKSW that we transferred onto a poly-D-lysine–coated 96 well plate (µ-Plate 96 Well Square ibiTreat, Ibidi, ref. 89626). The plate was then centrifuged at 200 × g for 1 minute at room temperature using a swinging-bucket rotor to allow the cells to gently adhere. Imaging was performed on a Nikon Ti2 spinning disk confocal system equipped with a Hamamatsu ORCA-Fusion Digital CMOS C14440-20UP camera. Z-stacks (0.2 µm intervals) were acquired using a Plan Apo λ 100× (NA 1.45) MRD01905 oil immersion objective.

All stacks were processed and analyzed in Fiji^79^. First, we performed a maximum Intensity Z-projection of the acquired Z-stacks; only the cells with their apico-basal axis parallel to the slide were retained for downstream analyses. We then manually drew the cell body contour using the *Polygon selections* tool, delimiting a region of interest from which the area, aspect ratio, and circularity were automatically extracted using the *Analyze>Measure* command. We measured the collar length by drawing a line with the *Straight Line* tool, starting at the base of the collar—the site directly connecting the microvilli to the cell body—and extending to the distal end of the microvilli. The length was automatically calculated using the *Analyze>Measure* command. The resulting data were plotted using GraphPad Prism 11.0.0.

### Immunostaining of actin and tubulin in gametes

To visualize the actin cytoskeleton in both fixed male and female gametes (Figure S7P), 200 μL of a starved culture diluted to 1 x 10^5^ cells/mL in AKSW (see “Choanoflagellate culturing”) was transferred to a poly-D-lysine–coated 96 well plate (µ-Plate 96 Well Square ibiTreat, Ibidi, Cat.No. 89626) and centrifuged at 200 × g for 1 minute at 22°C with a swinging-bucket rotor centrifuge to gently adhere the cells. The cells were then pre-treated with 6% Acetone diluted in AKSW at room temperature for 5 minutes without agitation, followed by fixation in 4% PFA in AKSW at room temperature for 15 minutes without agitation. After three washes with PEM (100 mM PIPES, pH 6.95; 2 mM EGTA; 1 mM MgCl2), the cells were permeabilized with the permeabilization/blocking buffer (1% [w/v] bovine serum albumin; 0.2% [v/v] Tween 20 in PEM) for 30 minutes at room temperature. Finally, the cells were incubated with 1 μg/mL rat anti-Tubulin antibody conjugated to Alexa Fluor 488 ([YOL1/34], Abcam, Cat. No. ab195883) in the permeabilization/blocking buffer for one hour at room temperature without agitation. After two washes with PEM, the cells were incubated with Rhodamine-labeled Phalloidin (Thermo Fisher Scientific, Cat. No. R415) diluted at 1:300 in PEM for 20 minutes at room temperature and imaged immediately using a Nikon Ti2 spinning disk confocal system equipped with a Hamamatsu ORCA-Fusion Digital CMOS C14440-20UP camera and a Plan Apo λ 100× (NA 1.45) MRD01905 oil immersion objective. Z-stacks were acquired at 0.2 µm intervals.

### Cupidon::mStayGold localization and dynamics

To assess the subcellular localization and dynamics of Cupidon (Figure 3C-J and 3M, Figure S7A-O and T, Videos S11,S12 and S15), we prepared cultures of Cupidon::mStayGold-expressing asexual cells or gametes at 5 x 10^5^ cells/mL into a final volume of 200 µL in AKSW that we transferred onto a poly-D-lysine–coated 96 well plate (µ-Plate 96 Well Square ibiTreat, Ibidi, ref. 89626). The plate was gently centrifuged at 200 × g for 1 minute at room temperature and directly imaged on a Nikon Ti2 spinning disk confocal system equipped with a Hamamatsu ORCA-Fusion Digital CMOS C14440-20UP camera. Single Z-plane time-lapses and Z-stacks were acquired using a Plan Apo λ 100× (NA 1.45) MRD01905 oil-immersion objective. Only the cells with their apico-basal axis parallel to the slide were kept for downstream analyses.

#### Measurement of Cupidon::mStayGold subcollar belt width

To compare the width of the Cupidon-enriched subcollar belt in both asexual cells and gametes (Figure 3F-G), Z-stacks of whole cells were acquired (0.2 µm intervals) under a 488nm laser with a 300 ms exposure time. We then performed a maximum intensity Z-projection of these stacks. A line spanning the width of the Cupidon::mStayGold subcollar belt was manually drawn along the apico-basal axis using the *Straight Line* tool in Fiji. The length of the lines, and therefore the subcollar belt widths, were then plotted using GraphPad Prism 11.0.0.

#### Measurement of Cupidon::mStayGold microdomain dynamics along microvilli

To capture the dynamics of Cupidon::mStayGold microdomains along the cell collar microvilli (Figure 3H and Figure S7N-O), we acquired a single Z-plane encompassing at least four full-length microvilli. Images were collected every 300 ms under a 488nm laser (with 300 ms exposure time, constraining the frame interval) over a total period of 10 s. The resulting time-lapses were then processed in Fiji as follows: first, we applied a background subtraction using the command Process > Subtract Background > *Rolling ball radius: 50 pixels* to increase the signal-to-noise ratio. Additionally, we enhanced contrast by running Process > Enhance Contrast > *Saturated pixels: 0.50% (Process all slices)*. We then tracked Cupidon microdomains using Manual Tracking in TrackMate^100^. The average speed for each track (defined here as the path traced by a single microdomain over at least three consecutive time points) was calculated, and the results were plotted using GraphPad Prism 11.0.0.

#### Measurement of Cupidon::mStayGold signal intensity along the microvilli

To assess the intensity of the Cupidon::mStayGold signal along the collar microvilli of asexual cells and female gametes (Figure 3I-J), Z-stacks were collected at 0.2 µm intervals under a 488nm laser with a 300 ms exposure time. We then performed a maximum intensity Z-projection of half of the collar (a subset of 15 Z-slices) from the original Z-stacks in Fiji. For each resulting projected image, we measured the mean background intensity value and subtracted it from the same image. All background-subtracted projected images were then combined into a single stack, and ROIs were manually drawn on each image. We used the *Straight Line* tool (Line Width: 3 pixels) to draw a line on the top of the microvilli, starting from the distal end of the microvilli (distance=0 µm) and extending at least 4µm towards the base of the microvilli. We then used the command *ROI manager>More>Multi Plot* to automatically extract pixel gray values along the microvilli axis (from 0 µ to 4µm). The resulting data were then plotted using GraphPad Prism 11.0.0.

### Live imaging of actin in male gametes

A starved culture of transgenic cells expressing Lifeact::mCherry (Lifeact::mCh)^101^ was prepared as described in the “*S. rosetta* culture conditions” section. A volume of 200 µL of culture was transferred onto an uncoated 96 well plate (µ-Plate 96 Well Square ibiTreat, Ibidi, ref. 89626). The plate was gently centrifuged at 200 × g for 1 minute at room temperature and directly imaged on a Nikon Ti2 spinning disk confocal system equipped with a Hamamatsu ORCA-Fusion Digital CMOS C14440-20UP camera. Z-stacks were acquired using a Plan Apo λ 100× (NA 1.45) MRD01905 oil-immersion objective.

### mRNA sequencing and transcriptomic analysis

Cultures were grown in triplicate for each culturing condition assessed. Samples of asexual cells were prepared from cultures of 1.5%/1.5% medium inoculated with 8,000 cells/mL of *S. rosetta* cells and supplemented with 167µg/mL of *E. pacifica* food bacteria in 3-layer culture flasks (Falcon™ Cell Culture Multi-Flask, 3-layer, 525 cm^2^, Corning, Cat. No. 89204-476). The cultures were incubated for 38 hours at 22°C and 60% humidity before collection. The same initial culture conditions were used to prepare starved cultures, with the difference that the cultures were incubated for 48 hours (early starvation), 60 hours (mid starvation), or 82 hours (late starvation) at 22°C and 60% humidity before collection. These time points were selected because they coincided with visible phenotypic transitions in the starving cultures, ranging from increased motility of asexual-looking cells during early starvation to fully differentiated male and female gametes during late starvation.

For each of the three replicates for each culturing condition (38, 48, 60, and 82 hours of incubation), 10^7^ cells were processed for lysis and RNA extraction. In practice, the cultures were pelleted and resuspended in AKSW, counted, and aliquoted to 1 x 10^7^ cells per aliquot. The aliquots were then centrifuged at 2800 × g for 10 minutes at 4 °C and resuspended in 100 μl of lysis buffer^102^: 20 mM Tris-HCl, pH 8.0; 150 mM KCl; 5 mM MgCl_2_; 250 mM sucrose; 1 mM DTT; 10 mM digitonin; 1 mg/mL sodium heparin; 1 mM Pefabloc SC; 100 μg/mL cycloheximide; 0.5 U/μl Turbo DNase (Thermo Fisher Scientific, Cat. No. AM2239); 1 U/μl SUPERaseIN (Thermo Fisher Scientific, Cat. No. AM2696) and incubated on ice for 10 minutes. Cells were passed ten times through a 30G needle and centrifuged at 6,000 × g for 10 min at 4°C. The supernatant was collected, brought to 100 μl with RNAse-free water, and RNA was purified using the RNAeasy Mini kit from Qiagen (Cat. No. 74104), eluting in 30 μl of RNase-free water.

RNA samples were measured with Qubit, and aliquots were prepared at 50 ng/mL in a final volume of 50 μl RNase-free water for each replicate and condition, and sent to SeqCoast Genomics (Portsmouth, NH, USA), who performed RNA library preparation and sequencing. Briefly, samples were prepared for whole-genome cDNA sequencing using an Illumina Stranded mRNA Prep (PolyA capture, ligation-based addition of adapters and indexes, see: https://www.illumina.com/products/by-type/sequencing-kits/library-prep-kits/stranded-mrna-prep.html) with IDT for Illumina Unique Dual Indexes. Sequencing was performed on the Illumina NextSeq2000 platform using a 200-cycle flow cell kit to collect between 23.2 million and 31.6 million 2×100 bp paired-end reads for each sample. Read demultiplexing, read trimming, and run analytics were performed using DRAGEN v3.10.12, the onboard analysis software on the NextSeq 2000. Reads were checked for quality by FastQC v0.12.0 (Babraham Bioinformatics), and we aligned them to predicted transcripts from the *S. rosetta* genome using Salmon v 1.5.2^103^. Transcript abundance and differential expression were quantified using edgeR^104^, both implemented within the DESeq2 v.1.42.0 software package^105^. TPM values for *cupidon* gene expression in asexual cells and early-, mid-, and late-starved cells, as well as differential expression tests comparing asexual cells to starved cells, are available in Figure S4A and Table S1. RNA sequencing data generated have been deposited to the NCBI Short Read Archive (NCBI BioProject Accession number: PRJNA1472704).

### CRISPR-Cas9 guide RNA and repair template design

We designed candidate CRISPR guide RNA (gRNA) sequences and repair templates for each gene of interest following previously established protocols^15^. Candidate CRISPR RNA (crRNA) sequences for the editing of *cupidon* were identified using the EuPaGDT tool^106^ and the *S. rosetta* genome. Guide RNA length was set to 15 nucleotides, and an expanded PAM consensus sequence, HNNRRVGGH, was used. The coding sequence for *cupidon* was obtained from the Ensembl Protists database (searching with the identifier PTSG_03655), which hosts the *S. rosetta* reference genome. crRNA candidates were filtered for guides with one on-target hit (including making sure the guides do not span exon-exon boundaries), zero off-target hits, lowest strength of the predicted secondary structure (assessed using the RNAfold web server: http://rna.tbi.univie.ac.at/cgi-bin/RNAWebSuite/RNAfold.cgi), and annealing near either the 5’ end of *cupidon* (*cupidon^1-^*^95^ and *cupidon^1–516^*) or the 3’ end of *cupidon* (*cupidon::ALFA* and *cupidon::mStayGold*), making sure to avoid conserved regions in the latter case by aligning with other choanoflagellate *cupidon* orthologs. crRNAs with the guide sequence of interest, as well as universal trans-activating CRISPR RNAs (tracrRNAs), were ordered from IDT (Integrated DNA Technologies) (File S4). Repair templates were designed as single-stranded DNA oligonucleotides matching the crRNA sense strand and containing homology arms corresponding to 50 bp of genomic sequence flanking each side of the Cas9-induced double-stranded break at the CRISPR target site. For gene truncation experiments (*cupidon^1-^*^95^ and *cupidon^1-^*^516^), the sequence 5’-TTTATTTAATTAAATAAA-3’, a “termination cassette” encoding a stop codon in every frame, was included between the homology arms. To ALFA-tag *cupidon*, a *S. rosetta* codon-optimized ALFA-tag sequence (5’-AGCCGCCTGGAGGAGGAGCTGCGCCGCCGCCTGACGGAG-3’)^51^ was inserted between homology arms. The interrupted codon at the cut site was restored, and we verified that the original reading frame was preserved (Figure S4B). Along with the crRNA targeting our gene of interest, we co-edited our cells using previously published sequences for a crRNA and repair template designed to introduce a P56Q mutation in *rpl36a*^15^, conferring cycloheximide resistance to enrich for genome-edited cells by selection. crRNA and repair oligo sequences are provided in File S4.

### CRISPR-Cas9 genome editing

Thirty-eight hours prior to the transfection, *S. rosetta* cells, either SrEpac or fluorescent transgenic lines, were seeded into 120 mL of 1.5%/1.5% culture medium at 8,000 cells/mL with 1 mL of *E. pacifica* (from a stock at 10 mg/mL *E. pacifica* suspension in ASW) in 3-layer culture flasks (Falcon™ Cell Culture Multi-Flask, 3-layer, 525 cm^2^, Corning, Cat. No. 89204-476) kept at 22°C and 60% humidity. This seeding density brings the culture to mid-log phase at the time of transfection. CRISPR/Cas9 nucleofection was performed according to the protocol outlined by Booth and King (2020)^15^ unless specified otherwise. Cultures were co-transfected with Cas9 RNPs and repair oligonucleotides targeting (1) the gene of interest and (2) the *rpl36a* locus to confer cycloheximide resistance (see “CRISPR guide RNA and repair template design”). The nucleofection reactions were prepared as described in Booth and King (2020)^15^, mixed into Lonza SF buffer (Lonza, Cat. No. V4SC-2960), added to a 96-well nucleofection plate (Lonza, Cat. No. V4SC-2960), and pulsed with the program CM156 using a Lonza 4D-Nucleofector (Cat. No. AAF-1003B for the core unit and AAF-1003S for the 96-well unit). Nucleofected cells were immediately recovered in recovery buffer (HEPES-KOH, pH 7.5; 0.9 M sorbitol; 8% [wt/vol] PEG 800) for 5-10 minutes and then added to 2 mL 1.5%/1.5% culture medium in a 6-well plate (Thermo Fisher Scientific, Cat. No. 140675) at 22°C and 60% humidity. After 30 min, 10 μL *E. pacifica* (from a stock at 10 mg/mL *E. pacifica* suspension in ASW) was added. Twenty-four hours later, 10 μL of 1 μg/mL cycloheximide (final concentration 5 ng/mL) was added to each well. Cycloheximide selection was carried out for five days. A negative control well of unedited cells (carried through the transfection experiment but without Cas9 RNPs or repair oligonucleotide) was used to confirm effective cycloheximide selection after the five-day selection period. To avoid starvation during this period, the nucleofected cultures were diluted on the third day at 1:50 into 2 mL of fresh 1.5%/1.5% culture medium supplemented with both *E. pacifica* and 10 μL of 1 μg/mL cycloheximide to maintain selection.

Following selection, the cells were diluted to 100,000 cells/mL and clonally isolated using a WOLF G2 Cell Sorter (NanoCellect Biomedical) equipped with a WOLF Microfluidic Sorting Cartridge (Cat. No. SP0511). Cells were sorted into 96-well plates (Thermo Fisher Scientific, Cat. No. 167008); each well contained 200 μL of 1.5%/1.5% culture medium supplemented with 50 μg/mL of *E. pacifica*. For each independent population of edited cells, between four and six 96-well plates were prepared. Cell sorting with this setup consistently yielded 70-80% of wells containing sorted cells per plate. Sorted cells were allowed to proliferate for 5 days to reach a workable cell density. To extract genomic DNA, 50 μL of cell culture was mixed with 50 μL of DNAzol direct (Molecular Research Center, Inc, Cat. No. DN131), incubated at RT for 10 minutes on a plate shaker (700 rpm), and stored at-20°C.

Genotyping PCRs were performed in 96-well plates (Brooks Life Sciences, Cat. No. 4ti-0770/c) using Q5 polymerase (NEB, Cat. No. M0491L) for 41 cycles. 2.5 μL of genomic DNA template was used in each 25 μL PCR reaction. PCR products were purified by magnetic bead cleanup and analyzed by Sanger sequencing (UC Berkeley DNA Sequencing Facility). Positive strains were expanded for downstream experimentation and long-term storage.

### Endogenous C-terminal fluorescent tagging of Cupidon

To visualize Cupidon protein in live cells, we used a recently developed protocol^60^, adapted from Combredet et al. ^14^ to generate a Cupidon::mStayGold strain. This technique enabled the in-frame insertion of a fluorescent marker at the 3’ end of the gene coding sequence to generate an endogenously tagged C-terminal fusion protein in *S. rosetta*. Briefly, we identified the most 3’-adjacent CRISPR/Cas9 PAM site within the *cupidon* coding sequence. We then engineered the pMS18 plasmid (Addgene ID 225681) so that it includes (1) the 59 bp region of the *cupidon* CDS downstream of the cut site (“*cupidon* 3’ CDS recovery region”), (2) a glycine-serine flexible linker sequence (SGGSGGS), fused in frame, (3) an in-frame codon-optimized *StayGold* coding sequence (gift from Jeffrey Colgren, Pawel Burkhardt lab, University of Bergen) ending with a stop codon, and (4) the 3’ UTR of *cupidon*, all positioned immediately upstream of the puromycin-resistance cassette. The resulting plasmid is available at Addgene (Addgene ID NK816; Cat. No. 257341).

PCR primers were then designed to amplify the repair cassette (*cupidon* 3’ CDS recovery region-linker - mStayGold coding sequence with a terminal stop codon - puromycin resistance cassette) with 50 bp homology arms framing the CRISPR/Cas9 cut site in the *cupidon* coding sequence. Information on the previously mentioned sequences is available in File S4. As described in Carver and King (2026)^60^, to ensure that no recombination between the two identical 50 bp sequences on the repair template – (1) the homology arm sequence of the 3’ primer, designed to match the genomic region adjacent to the cut site, and (2) the 3’ *cupidon* CDS recovery sequence encoded on the plasmid template – we introduced eleven synonymous nucleotide substitutions into the plasmid-encoded CDS recovery region to reduce similarity between these sequences while preserving the Cupidon amino acid sequence (File S4). The alternative codons were selected to match the original codon usage as closely as possible (File S4).

After PCR amplification and cleanup of the 3,343 bp repair template as outlined in Combredet et al. (2025)^14^, we recovered ∼15 μg of repair template in 30 μL nuclease-free water. We then transferred and evaporated 19 μL from this 30 μL volume (to achieve a final quantity of 10 μg of total repair template) at 55°C on a heating block for 2 h until 3μL were left in the tube and resuspended them directly in the final CRISPR reaction mixture (14 μL ice-cold SF buffer (Lonza, Basel, Switzerland; Cat. No. V4SC-2960), 4 μL SpCas9 RNP, 1 μg pUC19 (Nature Technology, Cat. No. NTC-RP500) carrier plasmid, and 2 μL of primed *S. rosetta* cells. A total of 50 reactions were performed. SpCas9 RNP and primed *S. rosetta* cells were prepared following the Booth and King (2020)^15^ protocol, as described in the “Genome editing” section. The use of a DG137 pulse^107^ was preferred over the CM156 pulse on the Lonza 4D-Nucleofector, as it appeared more effective, even though more cells lysed after the DG137 pulse in the SF Buffer. Transfected cells were then selected under 80 μg/mL puromycin treatment for 5 days at 22°C and 60% humidity. Puromycin-resistant cells were then screened for fluorescence on a Nikon Ti2 spinning-disk confocal system equipped with a Hamamatsu ORCA-Fusion Digital CMOS C14440-20UP camera. Fluorescent cultures were clonally isolated and genotyped by PCR and Sanger sequencing as described in “Genome editing” (primers listed in File S4).

### Generation of stable mCherry-and mStayGold-tagged membrane marker strains

Stable mCherry-and mStayGold::GG expressing *S. rosetta* strains were produced following a previously published method^101^ to design the plasmids, adapted from Wetzel et al. (2018)^46^, using the transfection protocol from Booth et al. 2018^102^. We generated the mCherry::GG and mStayGold::GG reporters by fusing a geranyl-geranylation (GG) sequence from the *S. rosetta* PTSG_00306 gene to the carboxy terminus of codon-optimized mCherry or mStayGold^52^ sequences. This strategy was first described by Booth et al. (2018)^102^, who demonstrated that transient expression of the mCherry::GG construct (Addgene ID NK644; Cat. No. 109096) successfully delineated cell outlines in *S. rosetta*. In practice, both mCherry::GG and mStayGold::GG were synthesized as gBlocks Gene Fragments (IDT), flanked by 20 bp homology arms to their digested destination vector, and then assembled in an NK802 vector (Addgene ID NK802; Cat. No. 166056) digested with *NcoI* and *AflII*, in a Gibson assembly reaction (NEB HiFi). In the resulting plasmids, expression of the mCherry::GG or mStayGold::GG construct is driven by an *S. rosetta* actin promoter and is flanked by a 3’UTR from *S. rosetta* elongation factor L (*EFL*) gene. The vector also carries the Pac puromycin resistance gene as an independent ORF, expressed under an EFL promoter^101^. These two plasmids were deposited on Addgene with the Addgene ID NK814(Cat. No. 257339) for mStayGold::GG and NK815 (Cat. No. 257340) for mCherry::GG.

Transfection of these plasmids in *S. rosetta* was performed according to Booth and King (2020)^15^, using 10 μg of plasmid in the final nucleofection mix, for a total of 50 reactions per plasmid (NK814 and NK815). The use of a DG137 pulse^107^ was preferred over the CM156 pulse on the Lonza 4D-Nucleofector, as it appeared more effective, even though more cells lysed after the DG137 pulse in the SF Buffer. Transfected cells were then selected under puromycin treatment for 5 days at 22°C and 60% humidity. Puromycin-resistant cells were then screened for fluorescence on a Nikon Ti2 spinning-disk confocal system equipped with a Hamamatsu ORCA-Fusion Digital CMOS C14440-20UP camera. Fluorescent cultures were clonally isolated and the derived cultures were expanded for downstream experimentation and long-term storage. We edited *cupidon* in both mCherry-and mStayGold::GG expressing *S. rosetta* strains (Figure 1I), following the same protocol described in the “CRISPR guide RNA and repair template design” and “Genome editing” sections.

### Microscopy and image acquisition

All DIC microscopy images and videos presented in this study were acquired on a Zeiss Axio Observer.Z1/7 Widefield microscope with a Hamamatsu Orca-Flash 4.0 LT CMOS Digital Camera (Hamamatsu Photonics) and either a Plan-Apochromat 10x/0.45 Ph 1 M27 objective with a 1.6x Optovar setting (Figures 1D and G, Figure 5F, Figures S2C-E, Videos S3, S4, S26), a Plan-Apochromat 20x/0.80 Ph 2 M27 with a 2.5x Optovar setting (Figures 2B and 2F, Figure 3L, Figure S7R-S, Video S5), a LD C-Apochromat 40x/1.1 Water Korr UV VIS IR with a 1.0x Optovar setting (Figure 1I, Figure 4A, Figure S8E, Figure S4E, Video S6, S8) or a 1.6x Optovar setting (Figure S7Q, Figure S8A Video S19), a Plan-Apochromat 100x/1.40 Oil DIC M27 with a 1.0x Optovar setting (Figure S2B, Figure S5A-D, Videos S2, S7).

All Fluorescence Spinning-Disk Confocal single Z-plane, Z-stacks (0.2 µm intervals), and Videos were acquired using a Nikon ECLIPSE Ti2 spinning-disk confocal system equipped with a Hamamatsu ORCA-Fusion Digital CMOS C14440-20UP camera and a Plan Apo λ 100× (NA 1.45) MRD01905 oil immersion objective. The Videos S9, S10, S13, S14, S16, S17, S20, S21, S23, S24, S25, S27, S28, S29, and S30 were further deconvolved using the ClearView-GPU™ Microscopy Deconvolution tool implemented in Imaris x64 v10.2.0.

## Statistics

Information about the quantification and statistical details of experiments can be found in the corresponding figure legends. Statistical tests and graphs were produced using GraphPad Prism 11.0.0.

## RESOURCE AVAILABILITY

### Lead contact

Further information and requests for resources and reagents should be directed to and will be fulfilled by the lead contact, Nicole King.

### Materials availability

Plasmids generated in this study have been deposited to Addgene (#NK814; Cat. No. 257339, #NK815; Cat. No. 257340, #NK816; Cat. No. 257341). Choanoflagellate cell lines used in this study are available upon request.

### Data and code availability

- RNA sequencing data generated in this study have been deposited to the NCBI Short Read Archive (NCBI BioProject Accession Number: PRJNA1472704).
- This paper does not report original code.
- Any additional information required to reanalyze the data reported in this paper is available from the lead contact upon request.

## ACKNOWLEDGEMENTS

We are grateful to Maxwell Coyle, Jules Lavalou, Chunzi Liu, and Pawel Burkhardt for their critical reading of the manuscript. We thank all current members of the King lab, as well as Maxwell Coyle, Thibaut Brunet, and David Booth, for valuable feedback throughout the project. We thank Flora Rutaganira for advice on western blots, Jeffrey Colgren for providing the *S. rosetta* codon-optimized mStayGold sequence, and Debbie Maizels for assistance with the figures. This work was supported by a Human Frontier Science Program long-term fellowship (LT 000919/2020-L) to A.G.D.L.B and an Investigator Award from the Howard Hughes Medical Institute (N.K.).

## AUTHOR CONTRIBUTIONS

Conceptualization, A.G.D.L.B. and N.K.; Formal analysis, A.G.D.L.B.; Funding acquisition, A.G.D.L.B. and N.K.; Investigation, A.G.D.L.B. and S.G.; Methodology, A.G.D.L.B. and N.K.; Supervision, N.K.; Visualization, A.G.D.L.B. and N.K.; Writing – original draft, A.G.D.L.B. and N.K.; Writing – review & editing, A.G.D.L.B. and N.K.

## DECLARATION OF GENERATIVE AI AND AI-ASSISTED TECHNOLOGIES

During the preparation of this work, the authors used Claude and ChatGPT to improve the manuscript’s readability and language and to double-check that figure references were correct. After using this tool, the authors reviewed and edited the content as needed and take full responsibility for the content of the published article.

## DATA AVAILABILITY

**Document S1. Figures S1-S9, Tables S1 and S2**

**Data S1. Supporting information for identification and multiple sequence alignment of Cupidon orthologs, MD and GAIN domains, genome editing, and cell fusion assay, related to Figures 1, 3, 4, and Figures S1, S5, and S8.**

**File S1. Cupidon and ortholog protein sequences.** FASTA file containing the *S. rosetta* Cupidon protein sequence and the sequences of its orthologs. Related to Figure S1A.

**File S2. Multiple sequence alignment of Cupidon and orthologs.** Full-length protein alignment of *S. rosetta* Cupidon and its orthologs. Related to Figure S1A.

**File S3. Mystery Domain (MD) protein sequences used for alignment and structural prediction.** FASTA file containing all MD protein sequences analyzed by multiple sequence alignment and structural prediction. Related to Figure S1C–E.

**File S4. gRNAs, repair templates, and PCR primers.** Sequences of all gRNAs, repair templates, and PCR primers used in this study. Related to Methods.

**File S5. C-terminal GAIN domain sequences.** FASTA file containing the C-terminal GAIN protein sequences used in the alignment. Related to Figure S5D.

**File S6. Density plots from cell fusion analyses.** All density plots collected from the analysis of cell fusion in replete and starved cultures. Related to Figure 4C and Figure S8B–D.

**Video S1.** Superimposed AlphaFold-predicted structures of Cupidon’s Mystery domain (red), with structural homologs found in Metazoa (orange), SAR (grey), Haptista (pink), Bacteria (green), and Viruses (blue), related to Figure S1E.

**Video S2.** High magnification of a clump observed in *cupidon*^1–95^ mutant culture, related to Figure 1D. Scale bar = 10 μm.

**Video S3.** Time-lapses of *cupidon^wt^*, *cupidon^1–516^*, and *cupidon*^1–95^ cultures after vortexing, related to Figure 1D. Scale bar = 50 μm.

**Video S4.** Rosette induction (+ RIF-OMVs) in both *cupidon^wt^* and *cupidon*^1–95^ cultures, related to Figure S2D. Scale bar = 50 μm.

**Video S5.** Z-stack of a clump encompassing a mixed population of mTFP-positive *cupidon^wt^* cells and mTFP-negative cupidon^1–95^ cells, related to Figure 1I. Scale bar = 30 μm.

**Video S6.** *cupidon^wt^* cultures under nutrient-replete, EroS_pv_ induced-, or starved conditions, related to Figure 2B. Scale bar = 30 μm.

**Video S7.** Latrunculin A disrupts the actin-based collar in asexual wild-type cells, abolishing collar structure compared to DMSO-treated controls, related to Figure 2F. Scale bar = 5 μm.

**Video S8.** *cupidon*^1–95^, *Cupidon::ALFA*, and *Cupidon::mStayGold* cultures, related to Figure S4E. Scale bar = 30 μm.

**Video S9.** 3D rendering of a live asexual cell expressing the membrane marker mStayGold::GG, related to Figure 3C. Scale bar = 5 μm.

**Video S10.** 3D rendering of a live asexual cell expressing Cupidon::mStayGold, related to Figure 3C’. Scale bar = 4 μm.

**Video S11.** Dynamics of Cupidon::mStayGold microdomains transported in both directions along the collar of an asexual cell, related to Figure 3H and Figure S7 N-O. Scale bar = 5 μm.

**Video S12.** An example of tracking of Cupidon::mStayGold microdomain, related to Figure S7 N-O. Scale bar = 1 μm.

**Video S13.** 3D rendering of a live female gamete expressing the membrane marker mStayGold::GG, related to Figure 3D. Scale bar = 3 μm.

**Video S14.** 3D rendering of a live female gamete expressing Cupidon::mStayGold, related to Figure 3D’. Scale bar = 5 μm.

**Video S15.** Dynamics of Cupidon::mStayGold microdomains transported in both directions along the collar of a female gamete, related to Figure S7 N-O. Scale bar = 5 μm.

**Video S16.** 3D rendering of a live male gamete expressing the membrane marker mStayGold::GG, related to Figure 3E. Scale bar = 5 μm.

**Video S17.** 3D rendering of a live male gamete expressing Cupidon::mStayGold, related to Figure 3E’. Scale bar = 2 μm.

**Video S18.** Dynamics of Cupidon::mStayGold in a male gamete, related to Figure 3E’. Scale bar = 3 μm.

**Video S19:** Example of mStayGold::GG-expressing male gamete (green) attached to a female gamete (DIC only), related to Figure 3L and Figure S7Q. Scale bar = 5 μm.

**Video S20.** Confocal spinning disk live microscopy of a male gamete attached to a female gamete via its Cupidon::mStayGold-enriched fertilopod, related to Figure 3M and Figure S7T. Scale bar = 5 μm.

**Video S21.** 3D rendering of a live pair of male and female gametes, attached via their respective Cupidon::mStayGold-enriched domains, related to Figure 3L and Figure S7Q. Scale bar = 4 μm.

**Video S22.** DIC and epifluorescence time-lapse of mStayGold::GG-expressing male and female *cupidon^wt^* gametes fusing together, related to Figure S8D. Scale bar = 5 μm.

**Video S23.** 3D rendering of a live asexual cell stained with the fluorescently labeled LEL lectin (cyan), related to Figure 5B and Figure S9 A-B. Scale bar = 2 μm.

**Video S24.** 3D rendering of a live asexual cell stained with fluorescently labeled Jacalin lectin (magenta), related to Figure 5D and Figure S9C and E. Scale bar = 2 μm.

**Video S25.** 3D rendering of a live Cupidon::mStayGold-expressing asexual cell stained with fluorescently labeled Jacalin (magenta), related to Figure 5D and Figure S9C. Scale bar = 3 μm.

**Video S26.** Untreated *cupidon*^1–95^, Jacalin-treated *cupidon*^1–95^, LEL-treated *cupidon*^1–95^, and STL-treated *cupidon*^1–95^ cultures of asexual cells, related to Figure 5F. Scale bar = 50 μm.

**Video S27.** 3D rendering of a live female gamete stained with the fluorescently labeled LEL lectin (cyan), related to Figure 5I and Figure S9F. Scale bar = 3 μm.

**Video S28.** 3D rendering of a live male gamete stained with the fluorescently labeled LEL lectin (cyan), related to Figure 5J and Figure S9H. Scale bar = 2 μm.

**Video S29.** 3D rendering of a live Cupidon::mStayGold-expressing male gamete stained with fluorescently labeled Jacalin (magenta), related to Figure S9J. Scale bar = 2 μm.

**Video S30.** 3D rendering of a live Cupidon::mStayGold-expressing female gamete stained with fluorescently labeled Jacalin (magenta), related to Figure S9J. Scale bar = 3 μm.

