## Supplementary Figures and Tables for "An adhesion GPCR regulates cell adhesion and mating in the closest living relatives of metazoans"

**This PDF file includes:**

Figures. S1 to S9
Tables S1 to S2

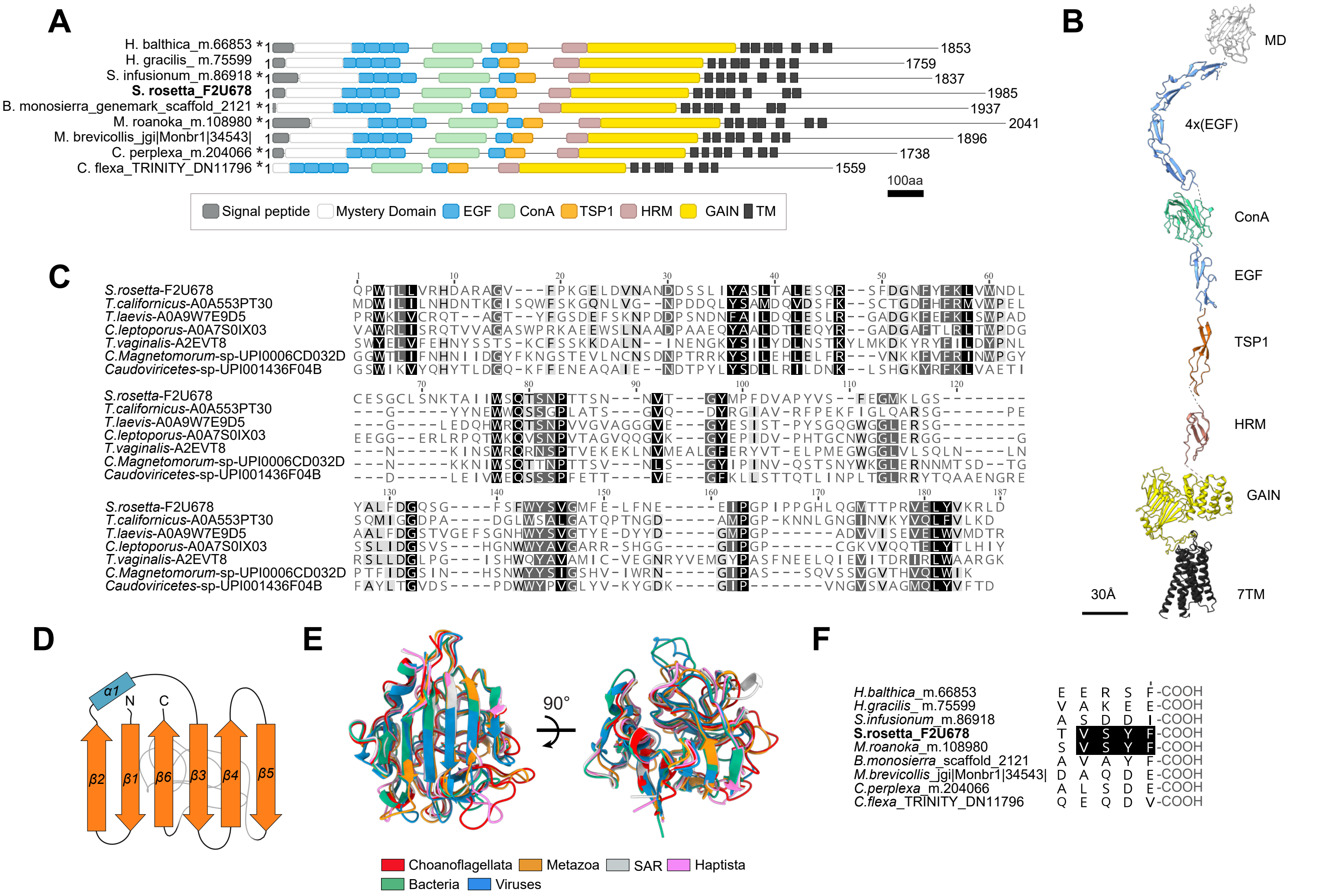
A
H. balthica_m.66853 *
H. gracilis_ m.75599
S. infusionum_m.86918 *
S. rosetta_F2U678
B. monosierra_genemark_scaffold_2121 *
M. roanoka_m.108980 *
M. brevicollis_jgi|Monbr1|34543|
C. perplexa_m.204066 *
C. flexa_TRINITY_DN11796 *
1853
1759
1837
1985
1937
2041
1896
1738
1559
100aa
Signal peptide
Mystery Domain
EGF
ConA
TSP1
HRM
GAIN
TM
B
MD
4x(EGF)
ConA
EGF
TSP1
HRM
GAIN
7TM
30Å
C
S.rosetta-F2U678
T.californicus-A0A553PT30
T.laevis-A0A9W7E9D5
C.leptoporus-A0A7S0IX03
T.vaginalis-A2EVT8
C.Magnetomorum-sp-UPI0006CD032D
Caudoviricetes-sp-UPI001436F04B
D
α1
N
C
β2
β1
β6
β3
β4
β5
E
90°
Choanoflagellata
Metazoa
SAR
Haptista
Bacteria
Viruses
F
H.balthica_m.66853 E E R S F -COOH
H.gracilis_m.75599 V A K E E -COOH
S.infusionum_m.86918 A S D D I -COOH
S.rosetta_F2U678 T V S Y F -COOH
M.roanoka_m.108980 S V S Y F -COOH
B.monosierra_scaffold_2121 A V A Y F -COOH
M.brevicollis_jgi|Monbr1|34543| D A Q D E -COOH
C.perplexa_m.204066 A L S D E -COOH
C.flexa_TRINITY_DN11796 Q E Q D V -COOH



Figure S1

**Figure S1. Cupidon protein domain architecture, structural conservation of the Mystery Domain, and identification of a putative PDZ-binding motif.** (A) The protein domain composition, order, and number of domains are largely conserved across Cupidon homologs from *S. rosetta* (in bold) and its closest choanoflagellate relatives. Protein domain identity is indicated in the key below the alignment. Asterisks denote N-terminal truncations due to incomplete sequence assemblies at the 5′ end of transcriptome-derived sequences. Scale bar, 100 amino acids. (B) *S. rosetta* Cupidon is a large, multidomain protein, as illustrated in this AlphaFold-predicted model of its extracellular region and 7TM domain. The hinges between domains and the long, disordered intracellular region of Cupidon, both of which are poorly resolved, have been removed (dashed lines). The domains were artificially aligned along a common axis for clarity; the relative orientation of the domains was adjusted for visualization purposes only. All structural models of the domains depicted both have a confidence score >70 Plddt. (C) Despite not having been previously identified as a distinct domain, the Mystery Domain (MD) sequence is conserved across phylogenetically distant organisms — including eukaryotes, bacteria, and viruses — suggesting it is under evolutionary constraint. Sequences shown are from representative eukaryotes (choanoflagellates, metazoans, SAR, metamonads, and Haptista), bacteria, and viruses. (D) The *S. rosetta* Cupidon Mystery Domain and its other eukaryotic, prokaryotic, and viral homologs adopt a β-strand-rich fold (β1→α1→β2→β6→β3→β4→β5) in which β6 is inserted between β2 and β3. A large, conserved loop separates β5 from β6. Schematic derived from the AlphaFold structural predictions. (E) The predicted 3D structure of the Mystery Domain is highly conserved across diverse eukaryotes, bacteria, and viruses, consistent with evolutionary constraint acting on this fold. View of the superimposed AlphaFold models from two different planes; color codes are indicated in the key. All structural models shown here have a confidence score >70 pLDDT. (F) *S. rosetta* and *M. roanoka* Cupidon share a predicted class I PDZ-binding motif (–VSYF, bold) at their C-termini, a feature not conserved across all choanoflagellate Cupidon homologs.

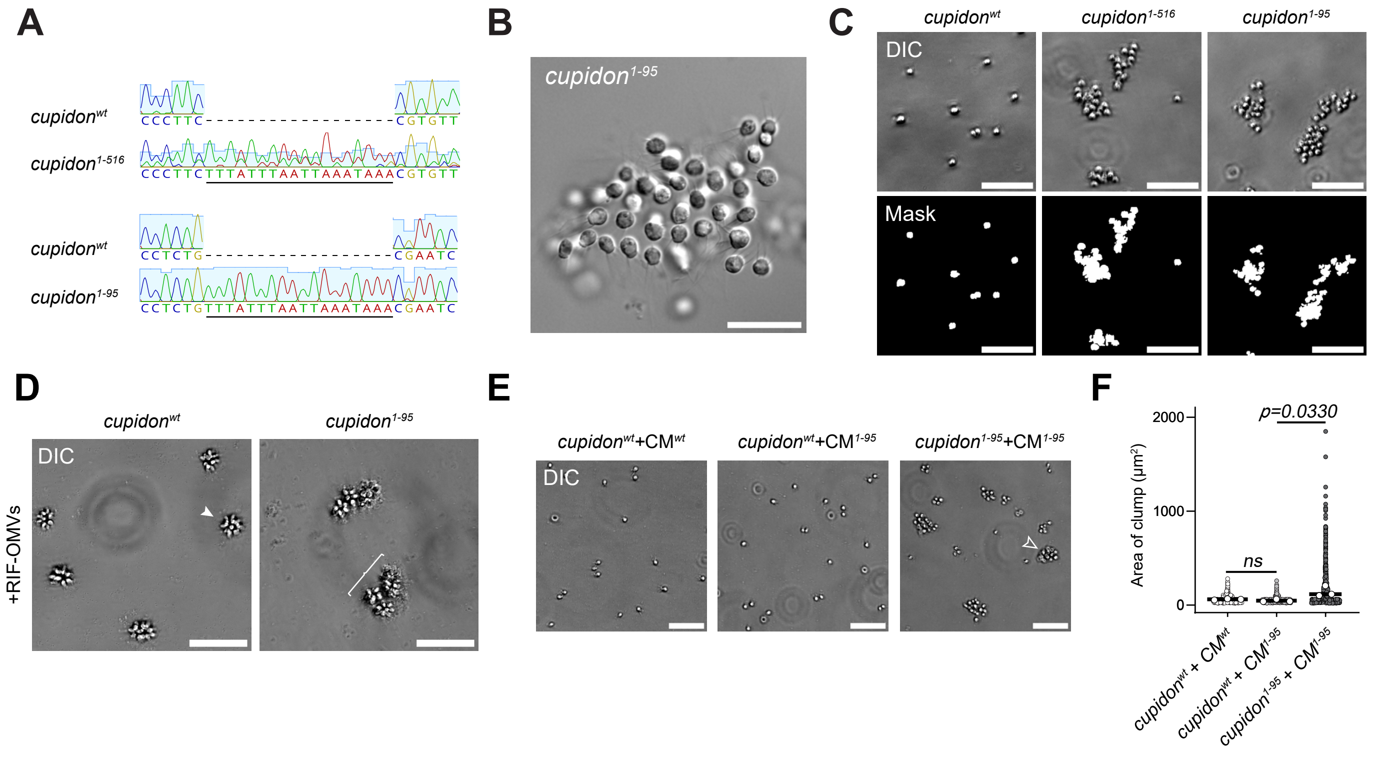
A
cupidon$^{wt}$
CCCTTC
cupidon$^{1-516}$
CCCTTCTTTATTTAATTAAATAAACGTGTT
cupidon$^{wt}$
CCTCTG
cupidon$^{1-95}$
CCTCTGTTTATTTAATTAAATAAACGAATC
B
cupidon$^{1-95}$
C
cupidon$^{wt}$
cupidon$^{1-516}$
cupidon$^{1-95}$
DIC
Mask
D
cupidon$^{wt}$
cupidon$^{1-95}$
DIC
+RIF-OMVs
E
cupidon$^{wt}$+CM$^{wt}$
cupidon$^{wt}$+CM$^{1-95}$
cupidon$^{1-95}$+CM$^{1-95}$
DIC
F
p=0.0330
ns
Area of clump (μm$^2$)
2000
1000
0
cupidon$^{wt}$ + CM$^{wt}$
cupidon$^{wt}$ + CM$^{1-95}$
cupidon$^{1-95}$ + CM$^{1-95}$



Figure S2

**Figure S2. Generation and characterization of *cupidon* mutant strains.** (A) CRISPR-mediated disruption of *cupidon* was confirmed by sequencing in both the *cupidon*$^{1-516}$ (top) and *cupidon*$^{1-95}$ (bottom) strains, each of which carries an early termination cassette[1] that truncates the Cupidon protein after amino acid 95 (*cupidon*$^{1-95}$) or 516 (*cupidon*$^{1-516}$). For each strain, the wild-type sequence is shown above for comparison. The inserted termination cassette is underlined. (B) *cupidon* mutant cells form large, disorganized clumps under nutrient-replete conditions, as shown in this representative DIC image. Scale bar, 20 µm. (C) Cell clumping was quantified using an automated image analysis pipeline. Representative DIC images of *cupidon*$^{wt}$, *cupidon*$^{1-516}$, and *cupidon*$^{1-95}$ cultures (top row) were converted into binary masks in which the projected area of each cell or clump was measured, yielding a measure of projected clump area as a proxy for the degree of cell aggregation (bottom row). This pipeline was used to quantify the data shown in Figure 1E and H, 2G, 5G, S2F, and S4F. Scale bar, 50 µm. (D) *cupidon*$^{1-95}$ cells form aberrant clumps of aggregated rosettes upon treatment with RIF-OMVs (bracket), whereas *cupidon*$^{wt}$ cells form isolated rosettes (white arrowhead). Representative DIC images are shown. Scale bar, 50 µm. See also Video S4. (E) The aggregation phenotype of *cupidon*$^{1-95}$ is cell-autonomous, as shown by conditioned medium (CM) transfer experiments. Cell clumping (empty arrowheads) was only observed in *cupidon*$^{1-95}$ cells treated with *cupidon*$^{1-95}$ CM, and not in *cupidon*$^{wt}$ cells treated with either *cupidon*$^{wt}$ CM or *cupidon*$^{1-95}$ CM. Representative DIC images are shown. Scale bar, 50 µm. (F) Quantification of CM transfer experiments confirmed that cell clumping is specific to *cupidon*$^{1-95}$ cells regardless of the source of CM. Projected clump area (µm²) is plotted for each condition. A minimum of 1372 projected areas (single cells or clumps) were plotted for each condition; 3 technical replicates were analyzed; white dots represent means per replicate; bars denote means across replicates. P-values were calculated from a one-way ANOVA with Tukey's multiple comparisons test.

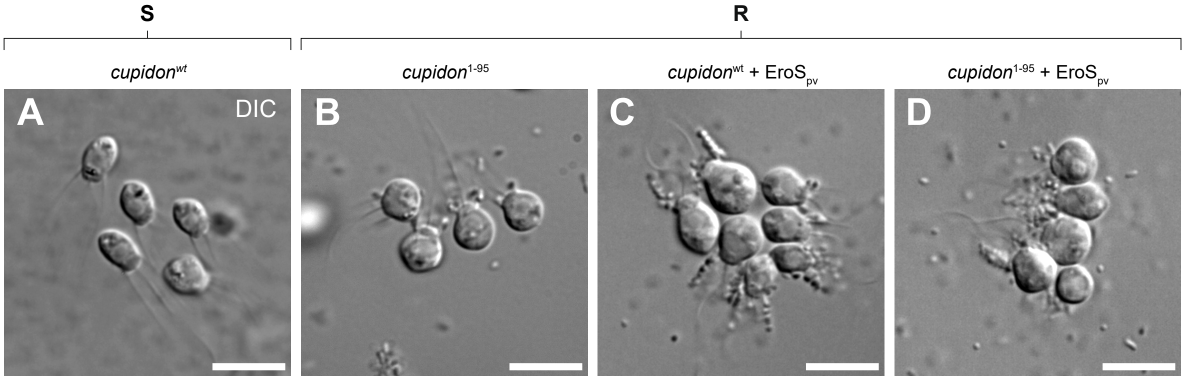

S
R
cupidonwt
cupidon1-95
cupidonwt + EroSpv
cupidon1-95 + EroSpv
A
DIC
B
C
D




Figure S3

**Figure S3. Cell-cell contact types differ between aggregates formed under spent medium and EroS-induced conditions.** Representative DIC images illustrating the predominant cell-cell contact types observed under four conditions: (A) *cupidon*$^{wt}$ in spent medium, (B) *cupidon*$^{1-95}$ in replete medium, (C) *cupidon*$^{wt}$ treated with $EroS_{pv}$ in replete medium, and (D) *cupidon*$^{1-95}$ treated with $EroS_{pv}$ in replete medium. In *cupidon*$^{wt}$ cells maintained in spent medium (A) and *cupidon*$^{1-95}$ cells in replete medium (B), collar-collar and cell body-collar contacts predominate (Fig. 2E). In contrast, treatment of *cupidon*$^{wt}$ cells with $EroS_{pv}$ (C) or *cupidon*$^{1-95}$ cells with $EroS_{pv}$ (D) shifts the predominant contact type to cell body–cell body interactions (Fig. 2E). Scale bar, 10 µm.

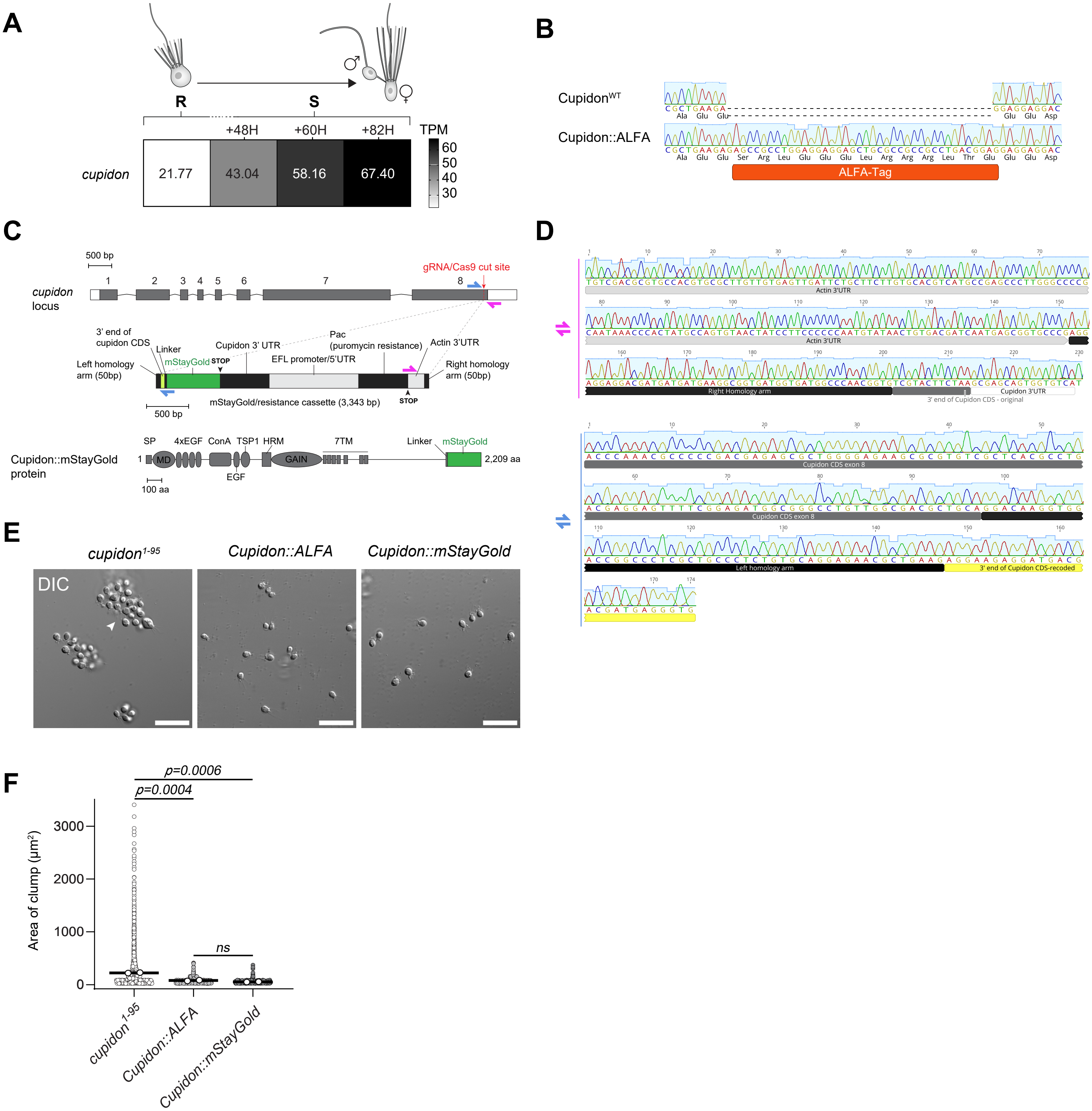
A
R
S
+48H
+60H
+82H
TPM
cupidon
21.77
43.04
58.16
67.40
60
50
40
30
B
Cupidon^WT
Cupidon::ALFA
ALFA-Tag
C
500 bp
cupidon locus
gRNA/Cas9 cut site
3' end of cupidon CDS
Linker
mStayGold
STOP
Cupidon 3' UTR
Pac (puromycin resistance)
Actin 3'UTR
EFL promoter/5'UTR
Left homology arm (50bp)
Right homology arm (50bp)
mStayGold/resistance cassette (3,343 bp)
Cupidon::mStayGold protein
SP
4xEGF
ConA
TSP1
HRM
7TM
Linker
MD
GAIN
EGF
2,209 aa
100 aa
D
Actin 3'UTR
Right Homology arm
Cupidon 3'UTR
3' end of Cupidon CDS - original
Cupidon CDS exon 8
Left homology arm
3' end of Cupidon CDS-recoded
E
cupidon^1-95
Cupidon::ALFA
Cupidon::mStayGold
DIC
F
p=0.0006
p=0.0004
ns
Area of clump (µm²)
3000
2000
1000
0
cupidon^1-95
Cupidon::ALFA
Cupidon::mStayGold



Figure S4

**Figure S4. *cupidon* is upregulated during sexual development, and endogenously tagged Cupidon fusion proteins retain wild-type anti-adhesive function.** (**A**) Expression of *cupidon* increases over the course of starvation. Transcripts per million (TPM) values for *cupidon* are shown for cells cultured under nutrient-replete conditions and at 48 h, 60 h, and 82 h after progressive depletion of the food bacteria in the culture medium. The schematic above illustrates the transition from asexual cells in replete conditions to differentiated male and female gametes after 82 h of growth in the same culture medium. Shading indicates average TPM value of *cupidon* across three biological replicates per condition. Results of the differential expression analysis using DESeq2 are provided in Table S1. **(B)** Sanger sequencing confirms in-frame insertion of the ALFA-tag coding sequence at the 3′ end of the endogenous *cupidon* gene in the Cupidon::ALFA strain. **(C)** Strategy for endogenous tagging of Cupidon with mStayGold. (Top) Intron/exon structure of the *cupidon* locus, comprising 8 exons (grey rectangles) and 7 introns. Target insertion site (red arrow). (Middle) The mStayGold/resistance cassette included 50 bp left and right homology arms, the recovered residual 3’ end of *cupidon* CDS left after the cut (yellow), a short linker, the mStayGold coding sequence (with stop codon), and the *cupidon* 3′ UTR, fused to the *S. rosetta efl* promoter and 5′ UTR driving expression of the puromycin resistance gene *pac*, followed by the *S. rosetta* actin 3′ UTR. PCR primers (magenta and blue half-arrows) were designed to verify the in-frame insertion of the cassette; the corresponding PCR products are presented in Figure S5D. (Bottom) The resulting locus encodes the full complement of Cupidon protein domains followed by mStayGold via a short linker. Scale bar, 100 aa. (**D**) Sanger sequencing confirms correct integration of the mStayGold cassette at the 3′ end of the endogenous *cupidon* locus. (**E**) Both Cupidon::ALFA and Cupidon::mStayGold strains display wild-type cell behavior under nutrient-replete conditions, with no detectable aggregation (as assessed by DIC microscopy), in contrast to the *cupidon*$^{1-95}$ strain. This demonstrates that neither fusion tag disrupts Cupidon's anti-adhesive function. Representative DIC images are shown. Scale bar, 30 µm. **(F)** Quantification of projected clump area (µm²) confirms that Cupidon::ALFA and Cupidon::mStayGold strains do not clump, while *cupidon*$^{1-95}$ cells do. A minimum of 2384 projected areas (single cells or clumps) were plotted for each condition; 2 technical replicates were analyzed; white dots represent means per replicate; bars denote means across replicates. P-values were calculated from a one-way ANOVA with Tukey’s multiple comparisons test.

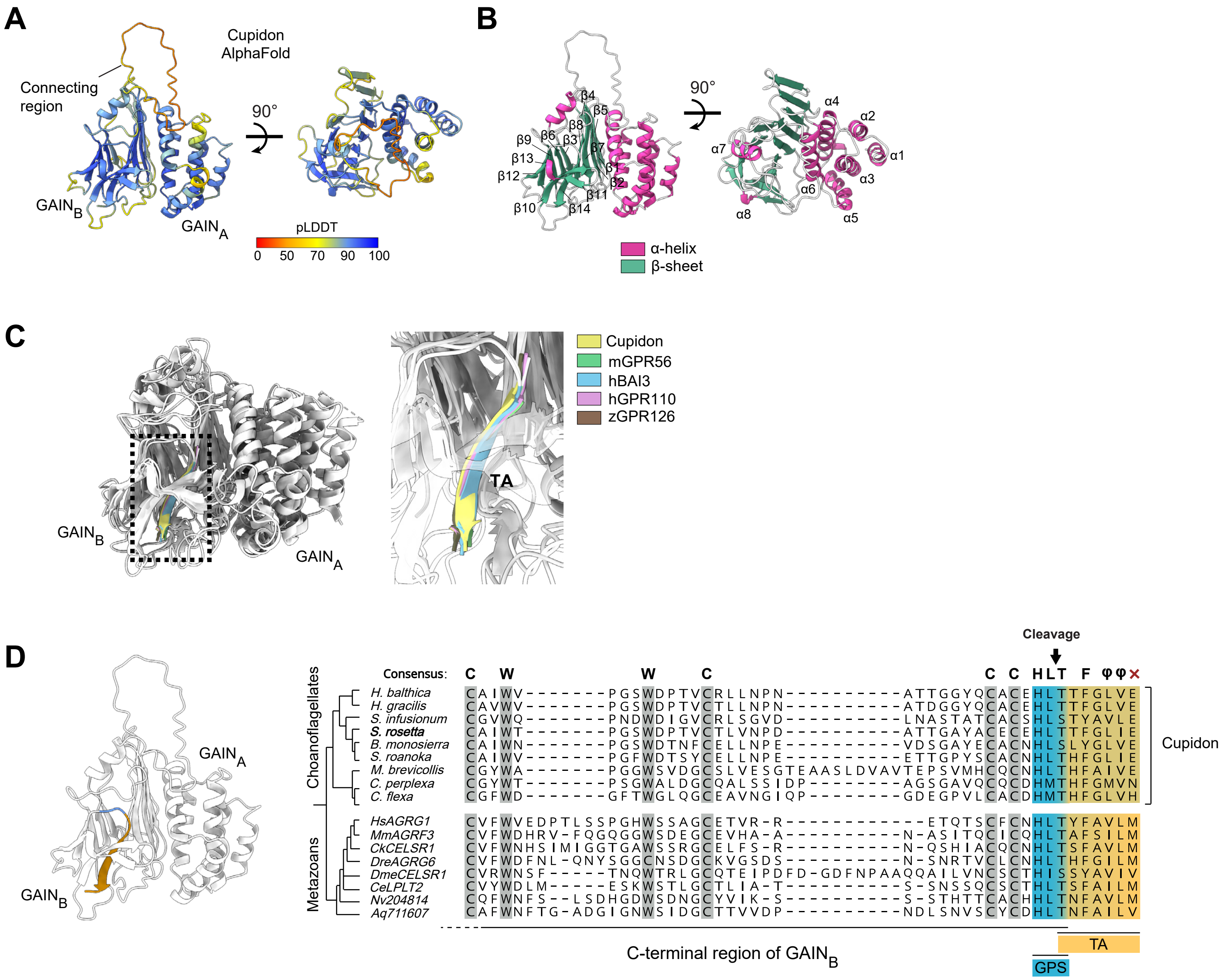

A
Connecting region
Cupidon AlphaFold
90°
GAINB
GAINA
pLDDT
0 50 70 90 100
B
β4 β5 β8 β6 β3 β9 β7 β13 β1 β12 β2 β11 β10 β14
90°
α4 α2 α7 α1 α3 α6 α8 α5
α-helix
β-sheet
C
GAINB
GAINA
TA
Cupidon
mGPR56
hBAI3
hGPR110
zGPR126
D
GAINA
GAINB
Cleavage
Consensus: C W W C C C HLT F φφ ×
Choanoflagellates
H. balthica CAIWV--------PGSWDPTVCRLLNPN-----------ATTGGYQCACEHLTTFGLVE
H. gracilis CAVWV--------PGSWDPTVCTLLNPN-----------ATDGGYQCACEHLTTFGLVE
S. infusionum CGVWQ--------PNDWDIGVCRLSGVD-----------LNASTATCACSHLSTYAVLE
S. rosetta CAIWT--------PGSWDPTVCTLVNPD-----------ATTGAYACECEHLTTFGLIE
B. monosierra CAIWN--------PGSWDTNFCELLNPE-----------VDSGAYECACNHLSLYGLVE
S. roanoka CAIWV--------PGFWDTSYCELLNPE-----------ETTGPYSCACNHLTHFGLIE
M. brevicollis CGYWA--------PGGWSVDGCSLVESGTEAASLDVAVTEPSVMHCQCNHLTHFAIVE
C. perplexa CGYWT--------PSGWALDGCQALSSIDP---------AGSGAVQCQCDHMTHFGMVN
C. flexa CGFWD--------GFTWGLQGCEAVNGIQP---------GDEGPVLCACDHMTHFGLVH
Cupidon
Metazoans
HsAGRG1 CVFWVEDPTLSSPGHWSSAGCETVR-R-------------ETQTSCFCNHLTYFAVLM
MmAGRF3 CVFWDHRV-FQGQGGWSDEGCEVHA-A-----------N-ASITQCICQHLTAFSILM
CkCELSR1 CVFWNHSIMIGGTGAWSSRGCELFS-R-----------N-QSHIACQCNHLTSFAVLM
DreAGRG6 CVFWDFNL-QNYSGGCNSDGCKVGSDS-----------N-SNRTVCLCNHLTHFGILM
DmeCELSR1 CVRWNSF-----TNQWTRLGCQTEIPDFD-GDFNPAAQQAILVNCSCTHISSYAVIV
CeLPLT2 CVYWDLM-----ESKWSTLGCTLIA-T-----------S-SNSSQCSCTHLTSFAILM
Nv204814 CQFWNFS--LSDHGDWSDNGCYVIK-S-----------S-STHTTCACHHLTNFAVLM
Aq711607 CAFWNFTG-ADGIGNWSIDGCTTVVDP-----------NDLSNVSCYCDHLTNFAILV
C-terminal region of GAINB
TA
GPS




Figure S5

**Figure S5. The Cupidon GAIN domain is structurally conserved with metazoan adhesion GPCRs and contains a predicted autoproteolytic cleavage site. (A)** The AlphaFold-predicted structure of the Cupidon GAIN domain consists of two subdomains ($\text{GAIN}_A$ and $\text{GAIN}_B$) connected by a flexible linker[2,3], colored according to the AlphaFold per-residue confidence score (pLDDT, red = low, yellow to blue = high). **(B)** The secondary structural elements of the Cupidon GAIN domain, with α-helices (pink) and β-sheets (green) colored and labeled. A total of 8 α-helices and 14 β-sheets are predicted in the GAIN domain of Cupidon. **(C)** The predicted Cupidon GAIN domain structure shows overall structural similarity with GAIN domains from metazoan aGPCRs, including the position of the Tethered Agonist (TA) within the $\text{GAIN}_B$ subdomain[3]; $\text{GAIN}_A$ and $\text{GAIN}_B$ subdomains are indicated. Inset: AlphaFold-predicted Cupidon GAIN domain structure and experimentally solved metazoan GAIN domain structures with color coding of the Cupidon TA (yellow) and the TA domains of four animal aGPCRs — mGPR56 (green), hBAI3 (blue), hGPR110 (pink), and zGPR126 (brown). The connecting region of Cupidon has been removed due to poor resolution. **(D)** The GPS autoproteolytic cleavage motif is conserved between choanoflagellate Cupidon orthologs and animal aGPCRs. (Left) AlphaFold-predicted structure of Cupidon with the C-terminal region of $\text{GAIN}_B$ highlighted, showing the GPS sequence (blue) and TA motif (yellow). (Right) Multiple sequence alignment of the C-terminal $\text{GAIN}_B$ region from choanoflagellate Cupidon homologs and representative animal aGPCRs. The predicted cleavage site is indicated by an arrow. The GPS sequence (blue) and TA motif (yellow) are highlighted. Conserved cysteine (C) and tryptophan (W) residues are also indicated above the alignment.

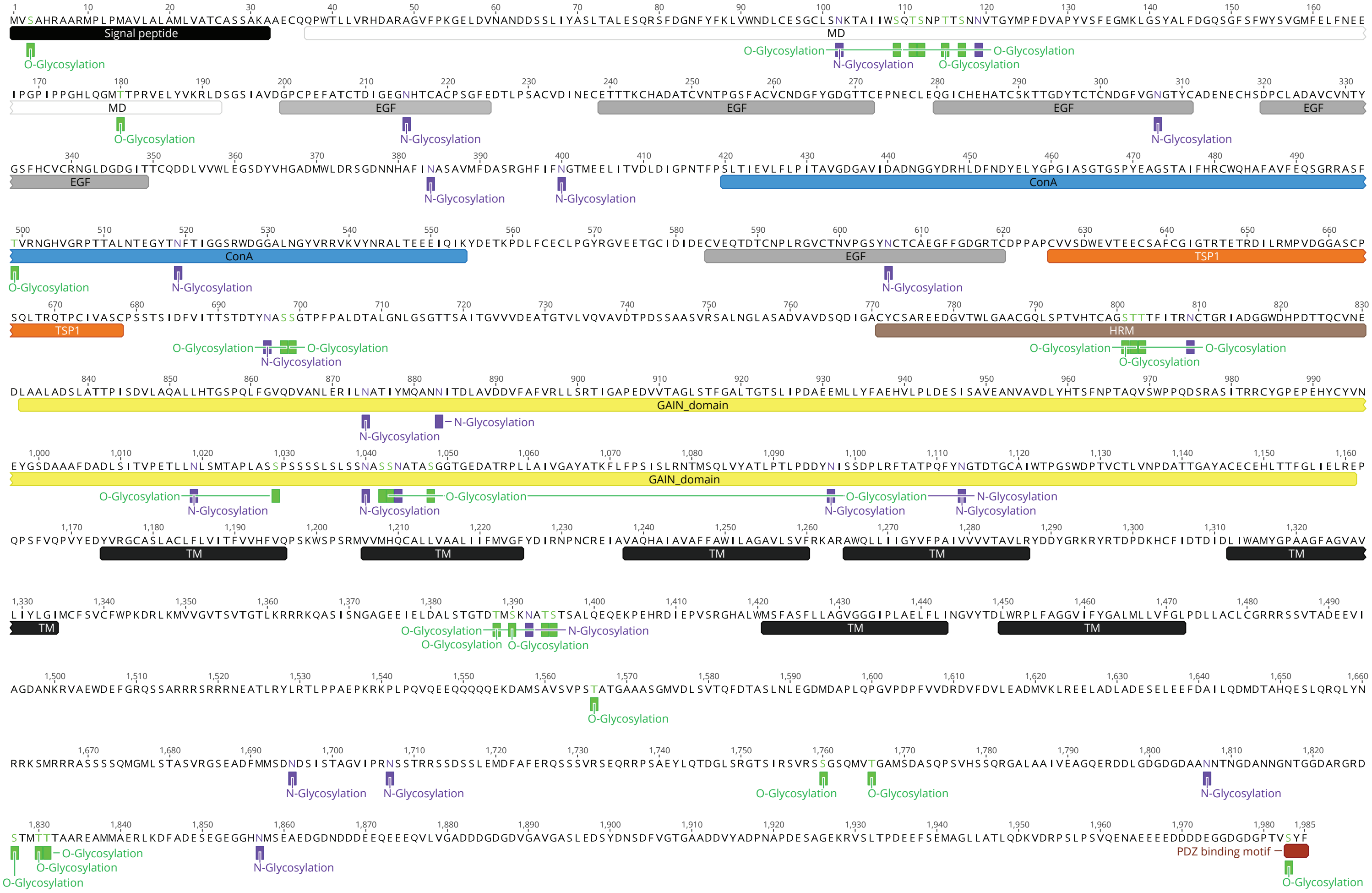
Signal peptide
MD
EGF
ConA
TSP1
HRM
GAIN_domain
TM
PDZ binding motif
O-Glycosylation
N-Glycosylation



Figure S6

**Figure S6. Predicted glycosylation of *S. rosetta* Cupidon.** The complete 1,985-amino acid sequence of *S. rosetta* Cupidon, annotated with predicted protein domains (colored rectangles, labeled) and predicted glycosylation sites. Predicted N-glycosylation sites are indicated in purple and predicted O-glycosylation sites in green. Domains are arranged in order from N- to C-terminus: Signal Peptide, Mystery Domain (MD), four EGF repeats, ConA, another EGF repeat, TSP1, HRM, GAIN, and seven transmembrane helices (TM1–7), followed by a predicted C-terminal PDZ-binding motif (–VSYF). N- and O-glycosylation sites are particularly dense across the extracellular MD, HRM, and GAIN domains as well as in the intracellular loop connecting TM5 to TM6. The visualization was generated using Geneious software.

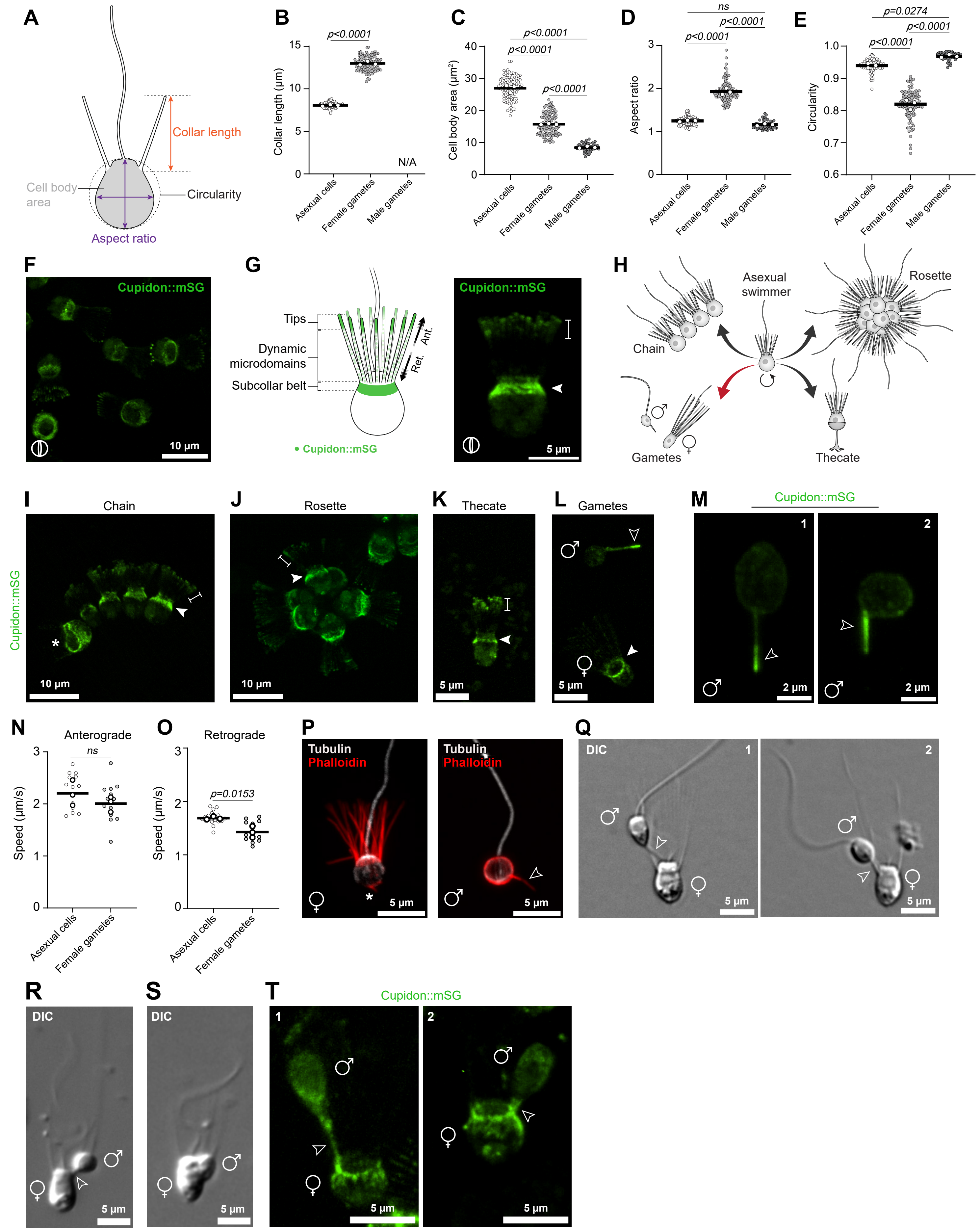
A
Collar length
Cell body area
Circularity
Aspect ratio
B
Collar length (µm)
p<0.0001
N/A
Asexual cells
Female gametes
Male gametes
C
Cell body area (µm²)
p<0.0001
p<0.0001
p<0.0001
D
Aspect ratio
ns
p<0.0001
p<0.0001
E
Circularity
p=0.0274
p<0.0001
p<0.0001
F
Cupidon::mSG
10 µm
G
Tips
Dynamic microdomains
Subcollar belt
Ant.
Ret.
• Cupidon::mSG
Cupidon::mSG
5 µm
H
Asexual swimmer
Chain
Rosette
Gametes
Thecate
I
Chain
Cupidon::mSG
10 µm
J
Rosette
10 µm
K
Thecate
5 µm
L
Gametes
5 µm
M
Cupidon::mSG
1
2
2 µm
2 µm
N
Anterograde
ns
Speed (µm/s)
Asexual cells
Female gametes
O
Retrograde
p=0.0153
Speed (µm/s)
Asexual cells
Female gametes
P
Tubulin
Phalloidin
5 µm
Tubulin
Phalloidin
5 µm
Q
DIC
1
2
5 µm
5 µm
R
DIC
5 µm
S
DIC
5 µm
T
Cupidon::mSG
1
2
5 µm
5 µm



Figure S7

**Figure S7. Morphological characterization of *S. rosetta* cell types and Cupidon::mStayGold localization across cell states. (A)** Schematic illustrating the morphological measurements used to quantify differences between asexual cells, female gametes, and male gametes. Collar length, cell body projected area, circularity, and aspect ratio (defined as the length along the apical–basal axis divided by the length along the equatorial axis of the cell body) are indicated. **(B)** Female gametes have longer collars (~13 µm on average) than asexual cells (~8 µm); male gametes entirely lack a collar (N/A). Three biological replicates were analyzed for each experimental condition; white dots represent means per replicate; bars denote means across replicates. A minimum of 100 cells per cell type were analyzed. P-values were calculated from an unpaired two-tailed t-test. **(C)** Asexual cells have the largest cell body projected area (~27 $\mu m^2$), followed by female gametes (~16 $\mu m^2$) and then male gametes (~8 $\mu m^2$). The projected cell body area ($\mu m^2$) is shown for all three cell types. Three biological replicates were analyzed for each experimental condition; white dots represent means per replicate; bars denote means across replicates. A total of 104 asexual cells, 111 female gametes, and 54 male gametes were analyzed. P-values were calculated from a one-way ANOVA with Tukey's multiple comparisons test. **(D–E)** Female gametes are elongated along the apical–basal axis relative to their equatorial diameter, with an aspect ratio around ~2 and reduced circularity (~0.8), whereas asexual cells and male gametes have a more circular morphology, with aspect ratios and circularity coefficients approaching 1 (~1.2 and ~0.94, respectively, for asexual cells; ~1.1 and ~0.97 for male gametes). Three biological replicates were analyzed for each experimental condition; white dots represent means per replicate; bars denote means across replicates. A total of 104 asexual cells, 111 female gametes, and 54 male gametes were analyzed. P-values were calculated from a one-way ANOVA with Tukey's multiple comparisons test. **(F)** A representative view of a culture of Cupidon::mStayGold-expressing asexual cells; Cupidon localizes to the collar microvilli and subcollar region. Scale bar, 10 µm. **(G)** (Left) Schematic of Cupidon::mStayGold distribution in asexual cells, illustrating enrichment at the microvillar tips and the subcollar belt, with anterograde and retrograde trafficking of Cupidon-containing microdomains along the microvilli indicated by arrows. (Right) Cupidon::mStayGold in a representative asexual cell, showing enrichment at the microvillar tips (bracket) and the subcollar region (arrowhead). Scale bar, 5 µm. **(H)** Schematic of select cell-state transitions in the *S. rosetta* life history. The asexual swimmer (center) can give rise to chains, rosettes, thecate cells, or differentiated gametes in response to different environmental cues. The transition to differentiated gametes (red arrow) is triggered by maintenance of cells in spent medium. **(I–K)** Cupidon::mStayGold is enriched at the microvillar tips (bracket) and subcollar region (arrowhead) in all examined asexual cell states, including chains (I), rosettes (J), and thecate cells (K). Scale bars, 10 µm (I, J), 5 µm (K). **(L)** Cupidon::mStayGold localizes to the subcollar region of female gametes (filled arrowhead) and to the fertilopod of male gametes (open arrowhead). Scale bar, 5 µm. **(M)** Cupidon::mStayGold distribution within the fertilopod varies between individual male gametes: enriched at the distal tip in cell 1 (open arrowhead) and distributed throughout the fertilopod in cell 2. Scale bar, 2 µm. **(N–O)** Anterograde (N) and retrograde (O) trafficking speeds of Cupidon::mStayGold microdomains along the microvilli are similar in asexual cells and female gametes (~2 µm/s and ~1.5 µm/s, respectively). Speed (µm/s) is plotted for each condition. Three biological replicates were analyzed per cell type, yielding a total of 17 asexual cells and 14 female gametes. Between 5 and 21 tracks, defined as the path traced by a single microdomain over at least 3 consecutive time points, were analyzed per cell. Each dot corresponds to the average speed of all anterograde or retrograde tracks for a given cell. P-values were calculated from an unpaired two-tailed t-test. **(P)** F-actin distribution differs markedly between female and male gametes. In female gametes (left), phalloidin staining is restricted to the apical collar microvilli and the basal cap of the cell (asterisk). In male gametes (right), F-actin occupies the entire cell cortex and extends into the fertilopod (arrowhead). Tubulin (white) and phalloidin (red) are shown. Scale bar, 5 μm. **(Q–S)** Representative DIC images of male–female gamete interactions. The fertilopod (open arrowhead) contacts the female gamete in the subcollar region (Q, R). Panel S shows two gametes caught in the act of cell fusion. Scale bars, 5 μm. **(T)** Cupidon::mStayGold imaging of two representative gamete pairs confirms that the Cupidon-rich fertilopod of the male gamete contacts the Cupidon::mStayGold-enriched subcollar zone of the female gamete. Open arrowheads indicate the fertilopod. Scale bar, 5 μm.

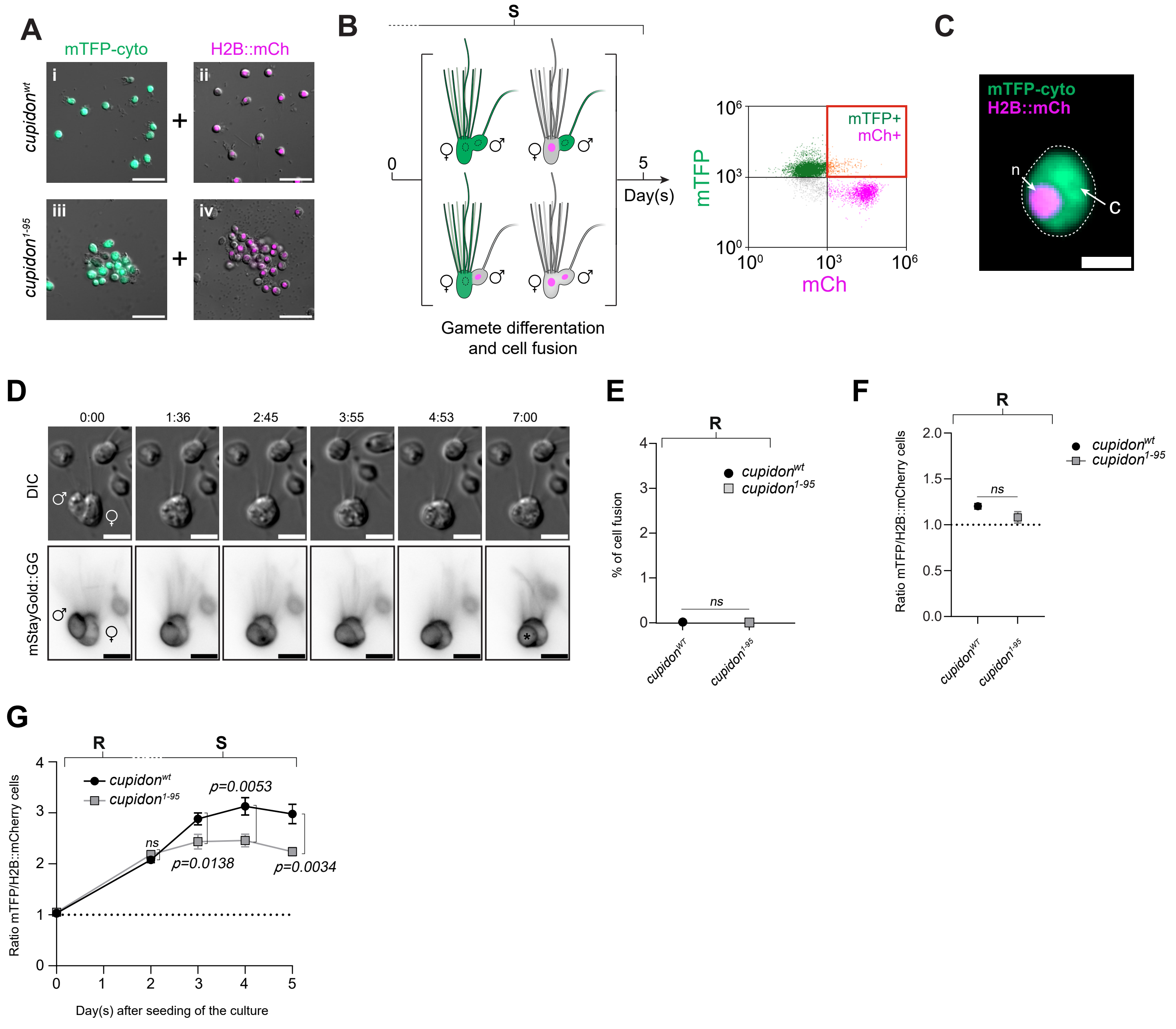
A
mTFP-cyto
H2B::mCh
cupidon^wt
cupidon^1-95
i
ii
iii
iv
+
B
S
0
5
Day(s)
Gamete differentation and cell fusion
mTFP
mCh
mTFP+
mCh+
10^6
10^3
10^0
C
mTFP-cyto
H2B::mCh
n
C
D
0:00
1:36
2:45
3:55
4:53
7:00
DIC
mStayGold::GG
E
R
% of cell fusion
cupidon^wt
cupidon^1-95
ns
cupidon^WT
cupidon^1-95
F
R
Ratio mTFP/H2B::mCherry cells
cupidon^wt
cupidon^1-95
ns
cupidon^WT
cupidon^1-95
G
R
S
cupidon^wt
cupidon^1-95
p=0.0053
ns
p=0.0138
p=0.0034
Ratio mTFP/H2B::mCherry cells
Day(s) after seeding of the culture



Figure S8

**Figure S8. *cupidon*$^{1-95}$ gametes have reduced cell fusion. (A)** Four different strains of cells were constructed to measure cell fusion efficiency in wild-type and *cupidon* mutant cells: *cupidon*$^{wt}$ expressing the cytoplasmic marker mTFP (mTFP-cyto), *cupidon*$^{wt}$ expressing the nuclear marker histone H2B fused to mCherry (H2B::mCh), *cupidon*$^{1-95}$ mTFP-cyto, and *cupidon*$^{1-95}$ H2B::mCh. Scale bars= 20 µm. **(B)** To measure cell fusion, we mixed *cupidon*$^{wt}$ mTFP-cyto cells with *cupidon*$^{wt}$ H2B::mCh cells or *cupidon*$^{1-95}$ mTFP-cyto cells with *cupidon*$^{1-95}$ H2B::mCh under starvation conditions. The percentage of cell fusion events was measured by flow cytometry at different time points after the initial mixing of tagged strains. An “event” (or cell) in which both mTFP and H2B::mCh were detected was scored as a cell fusion event. Assuming that fusion is random, this approach should detect half of the possible fusion events; fusion of cells with the same marker will not be detected. **(C)** An example of a fused cell observed in a starved mixed population of *cupidon*$^{wt}$ H2B::mCh and mTFP-cyto. The fluorescent signals of H2B::mCh and mTFP-cyto are detected within the same cell. Scale bar, 3 µm. **(D)** Timelapse of gamete fusion between a male (♂) and female (♀) gamete expressing the fluorescent membrane reporter mStayGold::GG (see Methods), shown in DIC (top) and the fluorescence (bottom) channels. Time is indicated in minutes:seconds. By the final time point (7:00), the male gamete has been entirely engulfed within the female gamete (asterisk). Scale bar, 5 µm. See Video S22. **(E)** Cell fusion does not occur under nutrient-replete conditions in either *cupidon*$^{wt}$ or *cupidon*$^{1-95}$ cultures. The average percentage of cell fusion events is plotted for each strain, with the standard deviation shown as error bars. A total of 30,000 events were analyzed from both wild-type and *cupidon*$^{1-95}$ mutant cultures, with three biological replicates per genotype. See also File S6. P-values were calculated from an unpaired two-tailed t-test. **(F)** After 20 hours of incubation, mTFP-cyto- and H2B::mCherry-labeled cells were still present at approximately equal proportions in both *cupidon*$^{wt}$ and *cupidon*$^{1-95}$ cultures (ratio ≈ 1, dashed line) grown under nutrient-replete conditions, verifying that the two experimental conditions can be directly compared. The average ratio and its standard deviation were calculated from three biological replicates, yielding a total of 30,000 events per condition. P-values were calculated from an unpaired two-tailed t-test. **(G)** Change in the ratio of mTFP- to H2B::mCherry-labeled cells under starvation in both *cupidon*$^{wt}$ and *cupidon*$^{1-95}$ cultures (related to Figure 4B). A drift in the ratio of mTFP- to H2B::mCherry-labeled cells was observed in both *cupidon*$^{wt}$ and *cupidon*$^{1-95}$ cultures over the five-day culturing period. By day 5, mTFP-positive cells outnumbered H2B::mCherry-labeled cells by 3-fold in *cupidon*$^{wt}$ cultures and 2.2-fold in *cupidon*$^{1-95}$ cultures. This imbalance likely caused an underestimation of total fusion events between mTFP- and H2B::mCherry-expressing cells in both experimental conditions. Importantly, however, this ratio drift does not account for the reduced fusion observed in *cupidon*$^{1-95}$ cultures. Indeed, the drift was less pronounced in *cupidon*$^{1-95}$ cultures than in *cupidon*$^{wt}$ cultures, yet gamete fusion was still reduced by approximately sevenfold in the mutant (Figure 4B). Thus, the labeling-ratio drift reinforces, rather than invalidates, the conclusion that *cupidon*$^{1-95}$ cells indeed exhibit a *bona fide* gamete fusion defect. The average ratios and their standard deviation were calculated from three biological replicates, yielding a total of 30,000 events per condition and per time point. P-values were calculated independently for each time point from a two-tailed t-test.

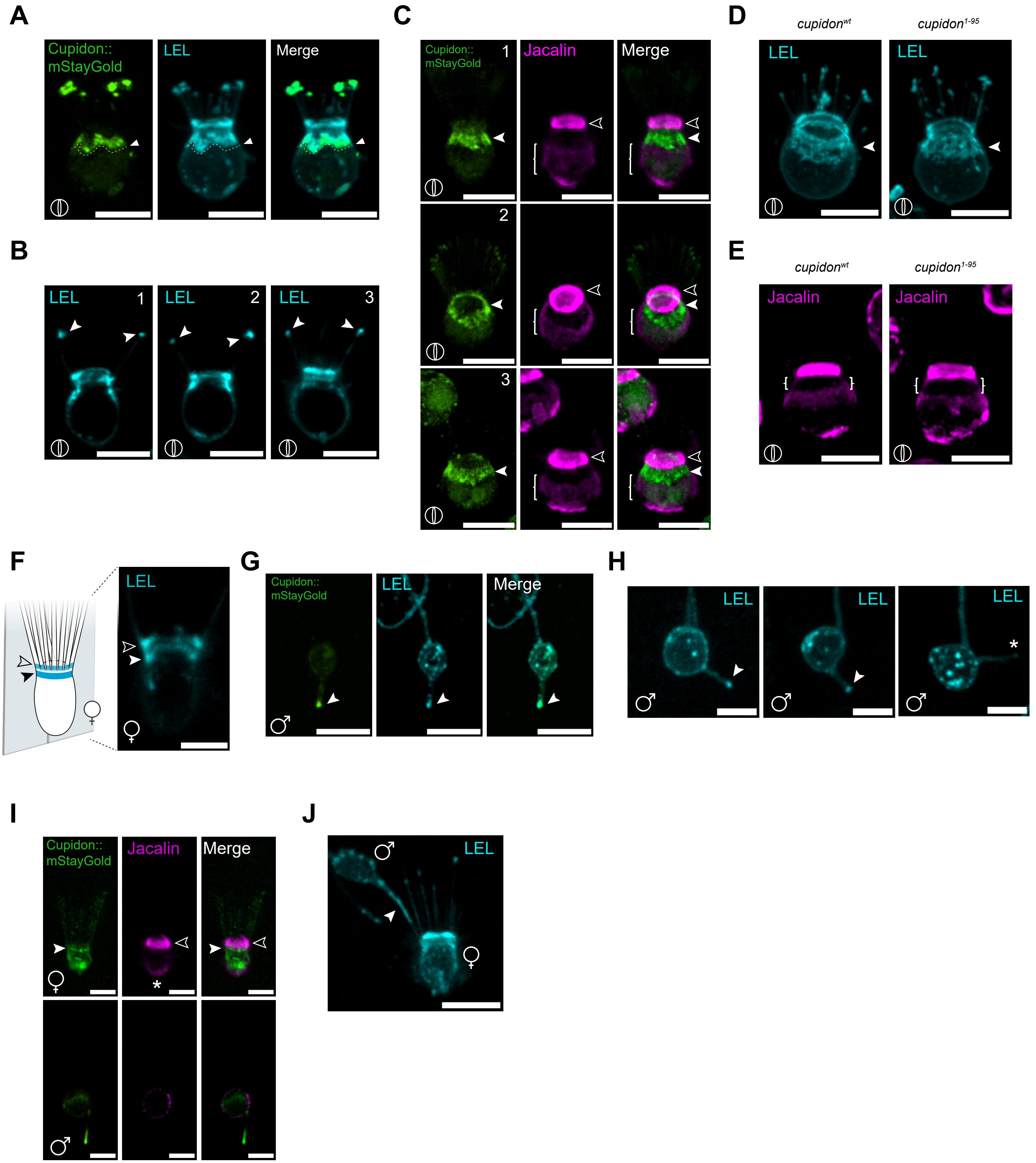
A
Cupidon::
mStayGold
LEL
Merge
B
LEL
1
2
3
C
Jacalin
D
cupidonwt
cupidon1-95
E
F
♀
G
♂
H
*
I
J



Figure S9

**Figure S9. N-acetylglucosamine-containing glycans are enriched at the collar of asexual cells and at the fertilopod of male gametes, independently of Cupidon. (A)** LEL staining colocalizes with Cupidon::mStayGold at the collar of asexual cells, with both signals enriched at the subcollar region and at the tip of microvilli (arrowhead). Scale bar, 5 µm. **(B)** Three representative single-Z-plane images of distinct LEL-stained asexual cells illustrate the distribution of LEL signal around the collar, including at the microvillar tips (arrowheads). Scale bar, 5 µm. **(C)** Jacalin staining and Cupidon::mStayGold signal are anti-correlated. Jacalin staining is specifically absent from the Cupidon-enriched subcollar belt and collar microvilli. A Jacalin-positive ring at the microvillar base (empty arrowhead) and a Jacalin-enriched lateral domain (bracket) flanked the Cupidon::mStayGold subcollar belt (filled arrowhead). Three representative cells are shown. Scale bar, 5 µm. **(D)** LEL staining is present at the collar and subcollar regions in both *cupidon*$^{wt}$ and *cupidon*$^{1\text{-}95}$ asexual cells (arrowhead), demonstrating that LEL-reactive glycans are displayed independently of Cupidon. Scale bar, 5 µm. **(E)** Jacalin staining is unaffected in *cupidon*$^{1\text{-}95}$ asexual cells compared to *cupidon*$^{wt}$ asexual cells (arrowhead), confirming that Jacalin-reactive glycans are also displayed independently of Cupidon. The position of the Cupidon-enriched subcollar belt is indicated (white brackets). Scale bar, 5 µm. **(F)** In female gametes, LEL staining is detected as two distinct rings: a narrow subcollar belt (filled arrowhead) and a second ring at the microvillar base (empty arrowhead), as illustrated in the schematic and corresponding cross-section fluorescence image. Only the LEL-positive narrow subcollar ring co-localizes with Cupidon in female gametes (see Figure 5I). Scale bar, 5 µm. **(G)** In male gametes, LEL staining colocalizes with Cupidon::mStayGold at the distal tip of the fertilopod (arrowhead). Scale bar, 5 µm. **(H)** Three representative male gametes showing LEL staining at the fertilopod (arrowhead). The absence of Cupidon::mStayGold signal at one fertilopod tip is indicated (asterisk). Scale bar, 3 µm. **(I)** Jacalin staining labels the ring at the microvillar base (empty arrowhead), and the basal cap (asterisk) of the female gamete. The Cupidon-enriched subcollar belt (filled arrowhead) does not overlap with Jacalin staining. In male gametes, Jacalin is weakly detected at the plasma membrane lining the cell body and is completely absent from the fertilopod, as shown in representative Cupidon::mStayGold, Jacalin, and merged images for female (top) and male (bottom) gametes. Scale bar, 3 µm. **(J)** During male–female gamete interactions, LEL staining marks the fertilopod of the male gamete as it contacts the female gamete (arrowhead). Scale bar, 5 µm.

1. Booth, D.S., and King, N. (2020). Genome editing enables reverse genetics of multicellular development in the choanoflagellate Salpingoeca rosetta. eLife *9*, e56193. https://doi.org/10.7554/eLife.56193.
2. Seufert, F., Pérez-Hernández, G., Pándy-Szekeres, G., Guixà-González, R., Langenhan, T., Gloriam, D.E., and Hildebrand, P.W. (2025). Generic residue numbering of the GAIN domain of adhesion GPCRs. Nat Commun *16*, 246. https://doi.org/10.1038/s41467-024-55466-6.
3. Langenhan, T., Anderson, G.R., Araç, D., Aust, G., Avila-Zozaya, M., Bagger, S.M., Barth, P., Berndt, S., Blacklow, S.C., Blanco-Redondo, B., et al. (2026). Adhesion G protein-coupled receptors. Pharmacological Reviews, 100116. https://doi.org/10.1016/j.pharmr.2026.100116.

**Table S1**. *cupidon* expression in starved cells compared with asexual cells.

| Gene | Life history stage | Log2 Fold change (vs asexual cells) | padj (adjusted P-value) | Interpretation |
|---|---|---|---|---|
| *cupidon* | Early starvation (+48H) | 1.078328 | 5.24E-62 | significantly different |
| | Mid starvation (+60H) | 1.157631 | 1.33E-80 | significantly different |
| | Late starvation (+82H) | 1.1822745 | 5.26E-68 | significantly different |

**Table S2**. Fluorescent lectins tested

| Lectins | Preferred Sugar Specificity* | Subcellular localization | | |
|---|---|---|---|---|
| | | *cupidon*$^{wt}$ asexual cells | *cupidon*$^{1-95}$ asexual cells | *cupidon*$^{wt}$ gametes |
| Con A (Concanavalin A) | Terminal α-Man3 and α-Man9, and biantennary N-glycans | Faint Collar ring | Faint Collar ring | Female gamete: Faint Collar ring; Male: n.d. |
| **Jacalin** | **GalNAcα, Core 1 and 3 O-glycans** | **Collar ring, Faint cell body-lateral domain, and Basal cap** | **Collar ring, Faint cell body-lateral domain, and Basal cap** | **Female: Collar ring, Faint and uniform cell body, and Basal cap; Male: Faint cell body** |
| **LEL (*Lycopersicon esculentum* (tomato) lectin)** | **GlcNAc oligomers, polyLacNAc, and/or chitin** | **Collar ring, Faint cell body, Subcollar belt, Tip of collar microvilli** | **Collar ring, Faint cell body, Subcollar belt, Tip of collar microvilli** | **Female: Collar ring, Faint Cell body, Subcolar belt, Faint whole microvilli; Male: Tip of the fertilopod, Faint cell body, and puncta at the cell membrane** |
| MAL I (*Maackia Amurensis* lectin I) | Terminal 3-O sulfated Gal on LacNAc, α2-3-sialylated LacNAc | Faint cell membrane | Faint cell membrane | Female and Male: Faint cell membrane |
| **STL (*Solanum tuberosum* (potato) lectin)** | **GlcNAc oligomers, polyLacNAc, and/or chitin** | **Collar ring, Faint cell body, Subcollar belt, Tip of collar microvilli** | **Collar ring, Faint cell body, Subcollar belt, Tip of collar microvilli** | **Female: Collar ring, Faint cell body, subcolar belt, Faint microvilli; Male: Tip of the fertilopod, Faint cell body, and puncta at the cell membrane cell body** |
| VVL (*Vicia villosa* agglutinin) | Terminal GalNAcβ, Terminal LacdiNAc | Some cells: Faint Collar ring | Some cells: Faint Collar ring | Some Female gametes: Faint collar ring; Male: n.d. |
| WGA (*Triticum vulgaris* (wheat germ) agglutinin) | GlcNAcβ, polyLacNAc, multiantennary N-glycans, chitin, terminal GlcNAcα-, GalNAcα-, GalNAcβ-, MurNAcβ- | Collar ring, Faint Cell body, Subcollar belt, Faint microvilli, Intermicrovillar bridges, and flagellar vanes | Collar ring, Faint Cell body, Subcollar belt, Faint microvilli, Intermicrovillar bridges, and flagellar vanes | Female: Collar ring, Faint Cell body, Subcollar belt, Faint microvilli, Intermicrovillar bridges, and flagellar vanes; Male: Tip of the fertilopod, Faint Cell body, and Puncta at the cell membrane |

**From* https://doi.org/10.1021/acschembio.1c00689 *and* Vector Laboratories Product Information

Symbols and abbreviations:

LacNAc: N-acetyllactosamine
GalNAc: N-acetylgalactosamine
GlcNAc: N-acetylglucosamine
Man: Mannose
n.d.: not detected
