## Supplementary figures and images for "An adhesion GPCR regulates cell adhesion and mating in the closest living relatives of metazoans"

### File S2

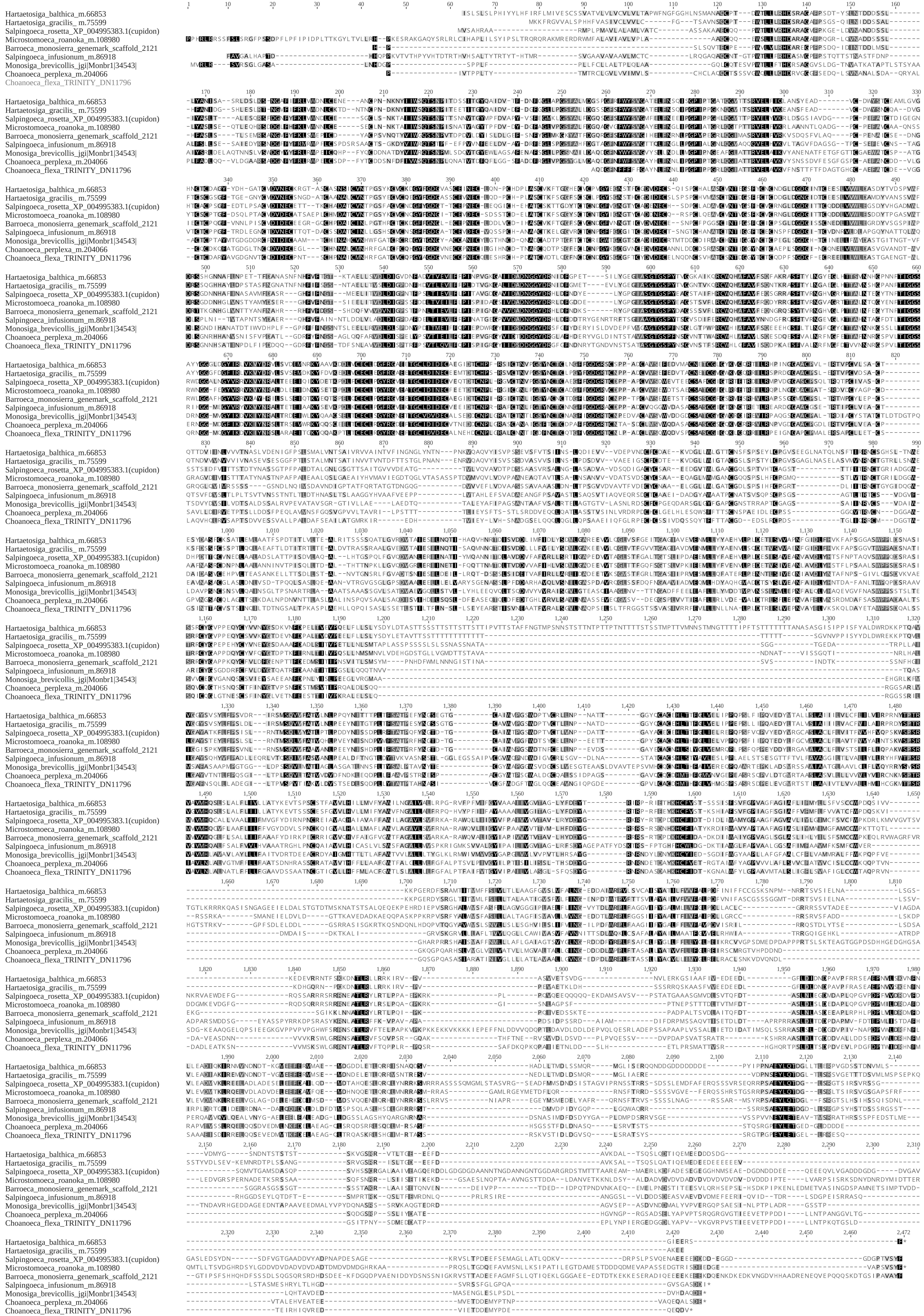

### File S6

#### Cell fusion assay_Nutrient-Replete (Day0, Day0+20H)

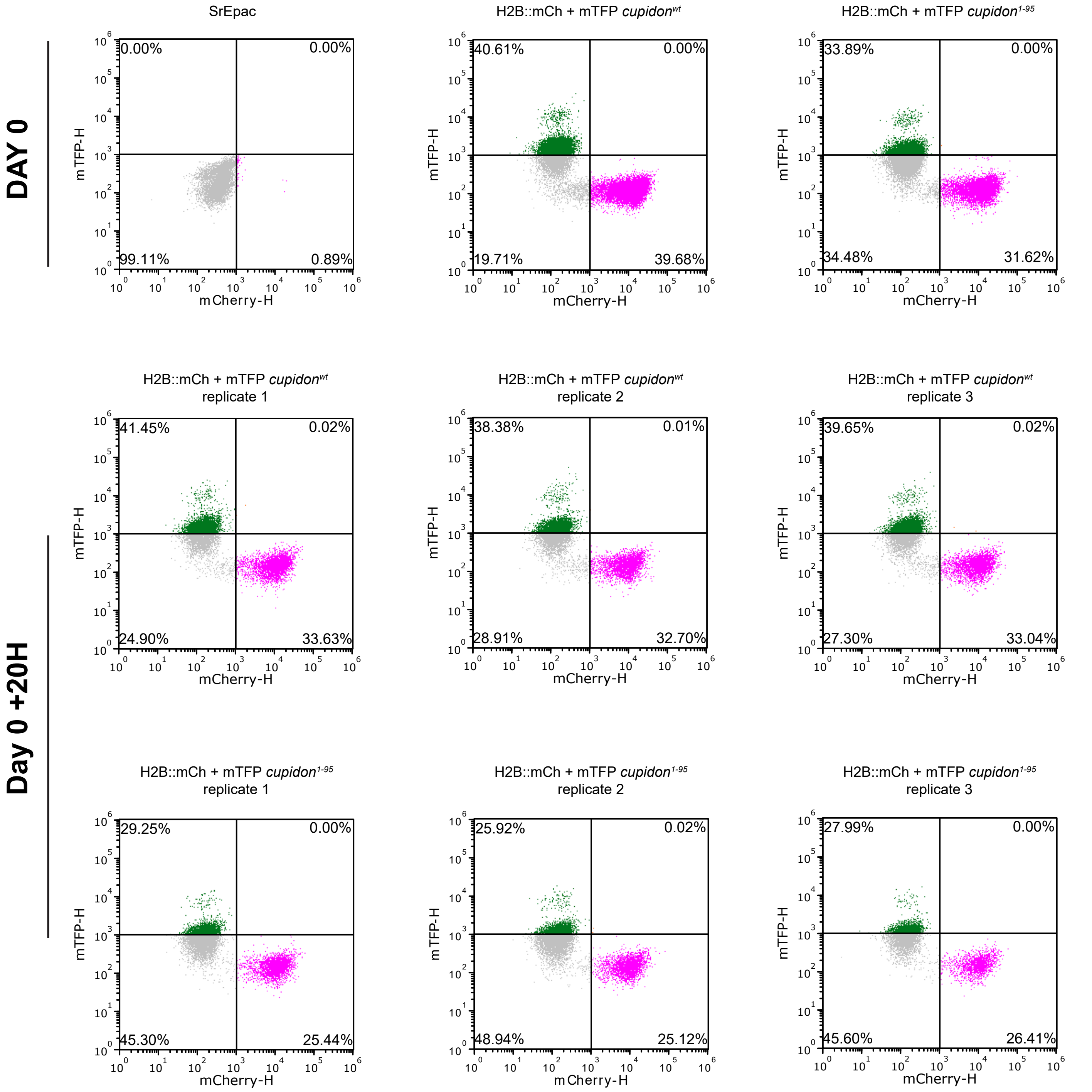

#### Cell fusion assay_Starvation (Day0-Day5)

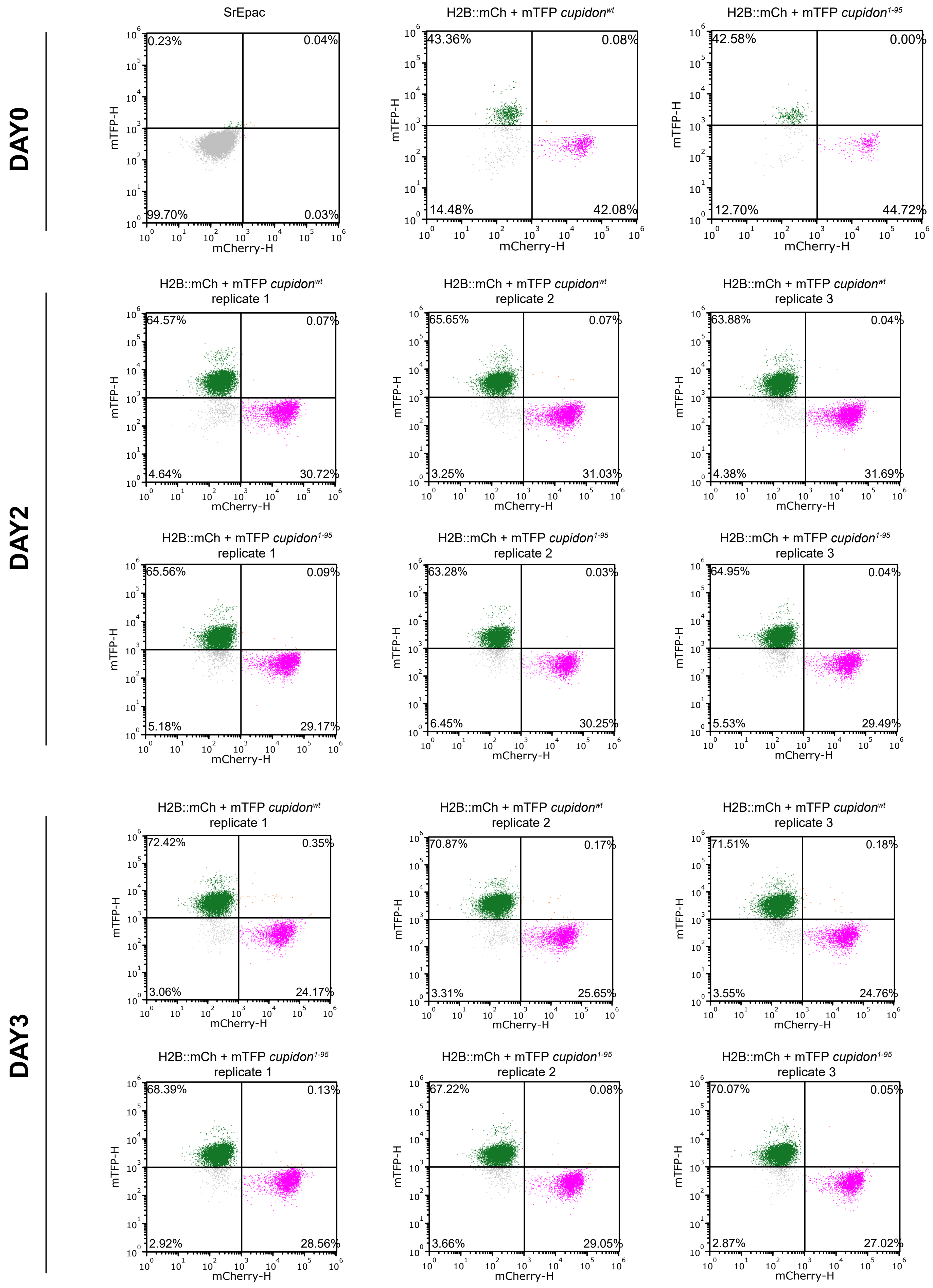

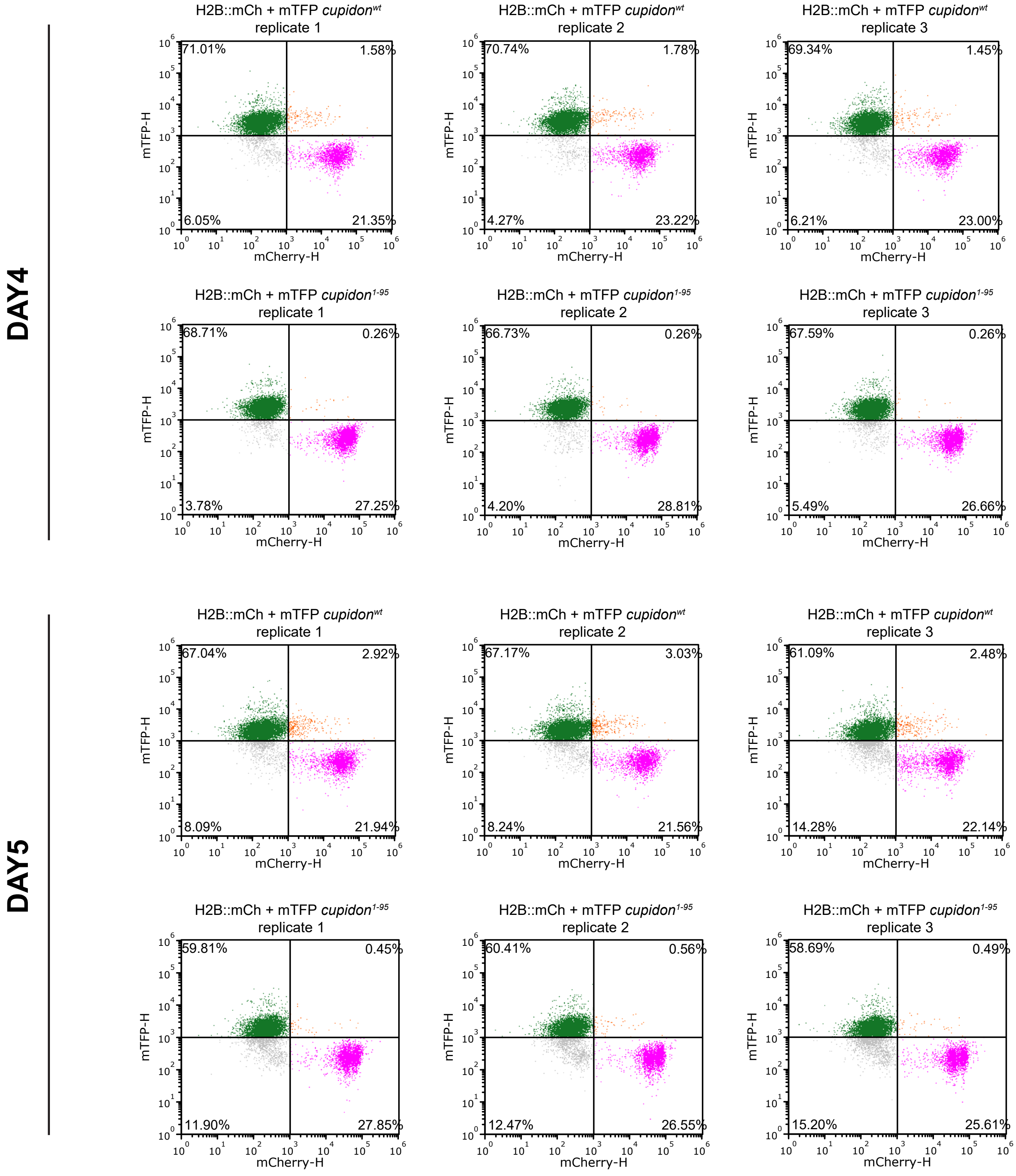
